# High-throughput transposon sequencing highlights the cell wall as an important barrier for osmotic stress in methicillin resistant *Staphylococcus aureus* and underlines a tailored response to different osmotic stressors

**DOI:** 10.1101/690511

**Authors:** Christopher F. Schuster, David M. Wiedemann, Freja C. M. Kirsebom, Marina Santiago, Suzanne Walker, Angelika Gründling

## Abstract

*Staphylococcus aureus* is an opportunistic pathogen that can cause soft tissue infections but is also a frequent cause of foodborne illnesses. One contributing factor for this food association is its high salt tolerance allowing this organism to survive commonly used food preservation methods. How this resistance is mediated is poorly understood, particularly during long-term exposure. In this study, we used TN-seq to understand how the responses to osmotic stressors differ. Our results revealed distinctly different long-term responses to NaCl, KCl and sucrose stresses. In addition, we identified the DUF2538 domain containing gene *SAUSA300_0957* (gene *957*) as essential under salt stress. Interestingly, a *957* mutant was less susceptible to oxacillin and showed increased peptidoglycan crosslinking. The salt sensitivity phenotype could be suppressed by amino acid substitutions in the transglycosylase domain of the penicillin binding protein Pbp2, and these changes restored the peptidoglycan crosslinking to WT levels. These results indicate that increased crosslinking of the peptidoglycan polymer can be detrimental and highlight a critical role of the bacterial cell wall for osmotic stress resistance. This study will serve as a starting point for future research on osmotic stress response and help develop better strategies to tackle foodborne staphylococcal infections.

## Introduction

*Staphylococcus aureus* is a Gram-positive bacterium that is carried by about 20 % of healthy individuals (Wertheim *et al*., 2005). It is also an opportunistic pathogen causing a variety of diseases, including bacteremia, endocarditis, soft tissue infections and food poisoning (Tong *et al*., 2015). One characteristic of *S. aureus* is its ability to grow in the presence of a high salt concentration (up to 2.5 M). Many other bacteria are unable to grow under these conditions (Measures, 1975), and this has been used extensively for the isolation of staphylococci. This salt tolerance also allows *S. aureus* to thrive in its niches such as the human nares, skin (Wertheim *et al*., 2005) and certain food products (Adams & Moss, 2008).

Bacterial cells are highly pressurized (up to 1.9 MPa for *Bacillus subtilis* (Whatmore & Reed, 1990)) and contain a strongly crowded cytoplasm that is hyperosmotic to the environment. Water is not transported into the cell actively but rather influxes passively into the cell due to the high internal osmolality. Whenever the extracellular osmolality increases, water exits the cell which can cause problems to multiple cellular processes, in part due to molecular crowding (Wood, 2011, van den Berg *et al*., 2017). Hence, *S. aureus* must possess strategies to counter the effects of salt damage, but these are not entirely understood, as most work has been performed in *Escherichia coli* and *B. subtilis*.

General mechanisms of hyper-osmotic stress mitigation in bacteria are the rapid import of potassium and the accumulation of small osmotically active compounds, named compatible solutes (reviewed in depth by (Bremer & Krämer, 2019, Gunde-Cimerman *et al*., 2018, Booth, 2014, Cox *et al*., 2018)). Potassium levels quickly subside in favor of an increase in the concentration of compatible solutes.

In *S. aureus*, results consistent with the uptake of potassium (Csonka, 1989, Csonka & Hanson, 1991, Sleator & Hill, 2002) followed by the accumulation of compatible solutes have been reported (Measures, 1975, Graham & Wilkinson, 1992, Christian & Waltho, 1964). The most effective compatible solutes in *S. aureus* are glycine betaine and proline (Miller *et al*., 1991, Measures, 1975, Townsend & Wilkinson, 1992, Anderson & Witter, 1982, Bae & Miller, 1992, Pourkomailian & Booth, 1992), which are imported by specific osmolyte transporters (Bae & Miller, 1992, Pourkomailian & Booth, 1992). These preferred osmolytes can accumulate in the cell to very high concentrations without negatively affecting cellular processes (Csonka & Hanson, 1991). In addition, *S. aureus* can also synthesize compatible solutes *de novo*, as in the case of glutamine (Anderson & Witter, 1982), but this process is much slower than the uptake of osmolytes and requires more energy.

Previous research investigating the underlying genetic factors of the *S. aureus* osmotic stress response implicated several osmolyte transporters in this process (Price-Whelan *et al*., 2013, Scybert *et al*., 2003, Schwan *et al*., 2006, Vijaranakul *et al*., 1998, Wengender & Miller, 1995). These include the potassium transport systems Kdp and Ktr (Price-Whelan *et al*., 2013), the proline transporter PutP (Wengender & Miller, 1995, Schwan *et al*., 2006), the arsenic transport system Ars (Scybert *et al*., 2003) and the branched chain amino acid uptake system BrnQ (Vijaranakul *et al*., 1998). In addition, although currently not experimentally verified in *S. aureus*, it can be assumed that the main glycine betaine uptake system OpuD (Wetzel *et al*., 2011, Zeden *et al*., 2018) is an important factor in *S. aureus* for osmotic adaptation as glycine betaine uptake reduces sensitivity to NaCl exposure (Miller *et al*., 1991). Of note, in recent time the second messenger cyclic-dinucleotide-AMP (c-di-AMP) (Corrigan & Gründling, 2013) has been tied to the regulation of osmotic stress in *S. aureus* and other Gram-positive bacteria (reviewed by Commichau *et al*., 2018). This is through the interaction and regulation of a number of potassium (Corrigan *et al*., 2013) and compatible solute transporters by c-di-AMP (Schuster *et al*., 2016).

Despite these multiple countermeasures, exposure of *S. aureus* to osmotic stress leads to morphological changes, including the formation of larger cells (Vijaranakul *et al*., 1995) and a thickening of the cell wall (Onyango *et al*., 2013). However, if and which genetic factors are required for those processes are currently not well understood. The cell wall of Gram-positive bacteria is an important barrier and acts as a counterpart to the pressurized cytoplasm. A major component of the cell wall is peptidoglycan, a polymer consisting of chains of repeating N-acetylglucosamine (GlcNAc) and N-acetylmuramic acid (MurNAc) units that are crosslinked with neighboring chains via short peptides and, in the case of *S. aureus*, pentaglycine cross-bridges (Reviewed by (Vollmer *et al*., 2008)). In the past, the contribution of the cell wall towards osmotic stress resistance was often not considered in the bacterial osmotic stress resistance field. Partly, because the integrity of the cell wall is often assumed to be a prerequisite rather than an active adaptation to survive osmotic stresses and partly because most cell envelope stress sensing systems are unable to directly sense osmotic stresses (Jordan *et al*., 2008).

Nevertheless, several possible changes to the cell wall have been observed in *S. aureus* during NaCl stress, such as shortened interpeptide bridges in the peptidoglycan (Vijaranakul *et al*., 1995), an increase in resistance to methicillin (Madiraju *et al*., 1987) and increased autolysis activity (Stapleton *et al*., 2007, Ochiai, 2000). In addition to peptidoglycan, other components within the cell envelope, such as teichoic acids and more specifically their modifications with D-alanine (Koprivnjak *et al*., 2006) and sugar residues (Mistretta *et al*., 2019, Kho & Meredith, 2018) are affected by osmotic stress. Combined, these findings indicate that the integrity of the cell wall is an important requirement for osmotic resistance.

Most studies on osmotic stress in *S. aureus* have been conducted using NaCl salt as the osmotic stressor. The ionic properties of NaCl have important implications on protein stability and activity as its accumulation can lead to denaturation of proteins. A few studies in *S. aureus* have also been performed with non-ionic osmotic stressors such as sucrose, sorbitol glycerol and amino acids (Stewart *et al*., 2005, Stewart *et al*., 2002, Mitchell & Moyle, 1959, Schwan *et al*., 2006). All of these different stresses are often used synonymously with osmotic stress, in part influenced by the findings that the initial mitigation of osmotic stress by either salts or sugars can be prevented by the accumulation of potassium and compatible solutes. However, there are potential differences in the long-term adaptation to ionic and non-ionic osmotic stressors, which up to date have not been investigated in detail. It is currently also not known if the adaptations besides the accumulation of potassium and osmolytes are similar or differ depending on the osmotic stressor, and this was addressed as part of this study.

Using a transposon insertion sequencing (TN-seq) method (Santiago *et al*., 2015), we determined on a whole genome level genes that are essential or dispensable during long-term osmotic stress caused by exposure to NaCl, KCl and sucrose. This yielded very different sets of essential and dispensable candidate genes, providing experimental evidence of the distinct nature of these stresses and linking a number of previously unknown factors to the osmotic stress response in *S. aureus*. Amongst other novel candidate genes, we identified gene *SAUSA300_0957* as an important NaCl resistance gene in *S. aureus* and show that the encoded protein is important for peptidoglycan homeostasis in *S. aureus*. Another protein, which we identified as important during salt stress, is the penicillin binding protein Pbp2, further emphasizing a key function of peptidoglycan during NaCl-induced osmotic stress. Taken together, these experiments highlight the bacterial cell wall as a key player in the salt stress response of *S. aureus*.

## Results

### Different types of osmotic stresses target different sets of genes

In the literature, osmotic stress is a loosely used term, however the type of ion or osmolyte can potentially have a great impact on how bacteria respond. To address this issue, we determined how the stress responses to different commonly used osmolytes compare. A highly saturated *S. aureus* transposon library was generated with a promoter-less transposon (Santiago *et al*., 2015) and subsequently used for transposon sequencing (TN-seq) experiments (Supplementary Table 1 for details on all TN-seq experiments). The library was propagated for 16 generations in either LB medium (Lennox) or in LB with extra 0.5 M NaCl, 0.5 M KCl or 1.0 M sucrose added (Fig. 1A). The molarity of sucrose was doubled to accommodate for the dissociation of the salts into two ions and hence leading to roughly the same osmolality. The culture challenged with 0.5 M NaCl grew the slowest, followed by 1.0 M sucrose, whereas the cells grown in 0.5 M KCl grew similarly to the cells grown in LB (Fig. 1B). As expected for a good quality transposon library, transposon insertions were found under all conditions in genes throughout the whole genome (Fig. 1C). Next, the number of TN-insertions per gene following growth under the NaCl, KCl or sucrose stress condition was compared to the number of TN-insertions per gene after growth in LB medium (Supplementary Table 2) and in this manner conditionally essential (Supplementary Table S3) and dispensable (Supplementary Table S4) genes identified. Fold-changes in transposon insertions per gene and q-values were plotted in volcano plots (Fig. 1D). From this, it was evident that under KCl stress the number of essential and dispensable genes was much lower than for NaCl or sucrose stress, indicating a much less severe effect of KCl on *S. aureus* cells. This was also reflected when inspecting the gene lists of the top 30 essential or dispensable genes (Supplementary Table S3 and S4) as in the KCl condition only 15 and 2 genes respectively met the requirements of q-value ≤ 0.05 and a fold-change of 5. To identify common genes between conditions, the fold-change stringency was relaxed to 2-fold and the overlap of genes was visualized in Venn diagrams (Fig. 1E, individual genes in Supplementary Table S5). Three genes, namely *SAUSA300_0425* (*USA300HOU_0457, mpsA (nuoF*)) coding for a cation transporter of the respiratory chain (Mayer *et al*., 2015), *SAUSA300_0750* (*USA300HOU_0796, whiA*) coding for a protein of unknown function and *SAUSA300_0846* (*USA300HOU_0903*) encoding a possible sodium:proton antiporter were essential in all conditions. In the case of dispensable genes, only one gene, *SAUSA300_1255* (*USA300HOU_1294, mprF/fmtC*), coding for a phosphatidylglycerol lysyltransferase involved in the defense against cationic microbial peptides, was identified in all conditions (Fig. 1E, Supplementary Table S5). To assess similarities and differences on a gene function and cellular pathway level, functional Voronoi maps were generated from the conditionally essential (Supplementary Fig. S1) and dispensable (Supplementary Fig. S2) genes regardless of their p-values. These images highlighted the essentiality of the wall teichoic acid through *tagO* and the *dlt* operon (Supplementary Fig. S1A) and the transporters AapA and MgtE in both NaCl (Supplementary Fig. S1A) and KCl (Supplementary Fig. S1B) but not in the sucrose stress condition (Supplementary Fig. S1C). When the dispensable genes were investigated (Supplementary Fig. S2), the number and type of genes in the NaCl condition differed considerably from KCl and sucrose stress. Under NaCl stress (Supplementary Fig. S2A), in addition to the penicillin binding protein *pbp2*, a large number of genes involved in respiration were found to be dispensable (Supplementary Fig. S2A). The number of dispensable genes in the KCl (Supplementary Fig. S2B) and sucrose (Supplementary Fig. S2C) conditions were considerably lower. Taken together, the overlaps between conditions were small, indicating distinct modes of actions for each osmotic stress.

**Fig. 1.**
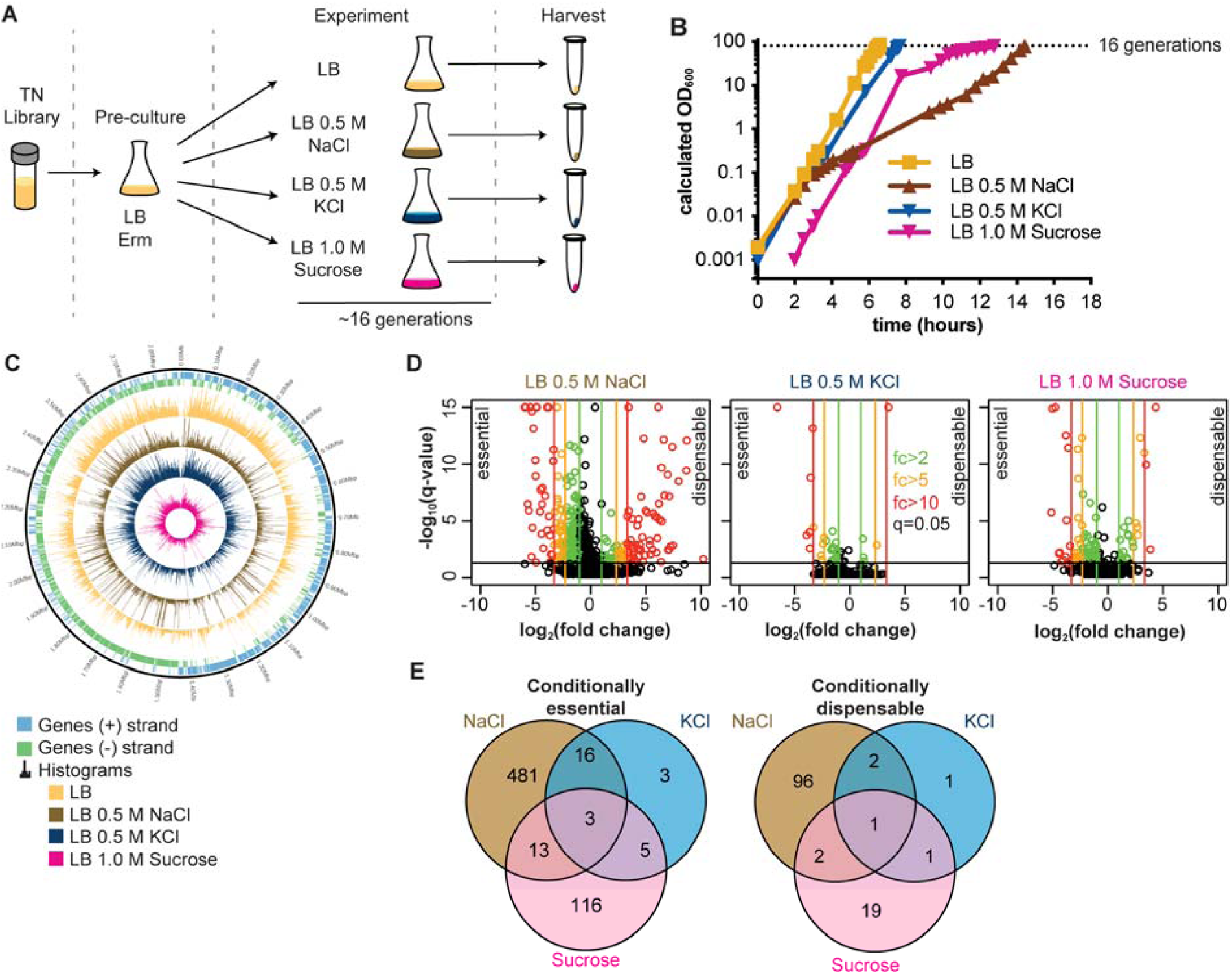
Different sets of genes are conditionally essential for the growth of *S. aureus* under NaCl, KCl or sucrose stress conditions. A. Workflow of the TN-seq experiment. A TN-seq library was pre-cultured and then used to inoculate LB, LB containing 0.5 M NaCl, 0.5 M KCl or 1 M sucrose. Cells were kept in exponential phase for 16 generations, harvested and transposon insertion sites determined by high throughput sequencing. B. Growth curves. For the TN-seq experiment (n=1), the growth of an *S. aureus* library cultured in LB, LB 0.5M NaCl, LB 0.5 M KCl or LB 1.0 M sucrose medium was followed by taking OD_600_ measurements at timed intervals. The cultures were back-diluted once when they reached an OD_600_ of approximately 0.3. The plotted OD_600_ values were calculated by multiplying the measured OD_600_ with the dilution factor. The dotted line indicates the 16 generation cut-off at which cultures were harvested. C. Circular plot showing the transposon insertion density along the *S. aureus* genome under different osmotic stress conditions. The two outer rings depict genes located on the (+) or (-) strand in *S. aureus* strain USA300 FPR3757. The inner four rings show the histograms of transposon insertions on a per gene basis after growth of the library in LB (orange), 0.5 M NaCl (brown), 0.5 M KCl (blue) and 1 M sucrose (pink) medium for 16 generations. D. Volcano plots of conditionally essential and dispensable genes following growth of *S. aureus* in 0.5 M NaCl, 0.5 M KCl or 1 M sucrose medium and compared to LB conditions. q-values represent Benjamini-Hochberg corrected p-values, and log_2_ (fold changes) indicate essential genes (negative values) or dispensable genes (positive values). The black horizontal line indicates a q-value of 0.05, which was deemed the significance level. Vertical lines indicate 2- (green), 5- (orange) or 10- (red) fold changes in either direction. Each dot represents one gene and coloring follows the fold- change scheme whenever the q-value threshold was met. Very small q-values were truncated to fit onto the graph. The TN-seq experiments were conducted once (n=1). E. Venn diagrams showing overlaps of conditionally essential and dispensable genes during prolonged NaCl, KCl or sucrose stress. Genes with a 2-fold decrease (essential genes) or a 2-fold increase (dispensable genes) in transposon insertions under each stress condition compared to the LB condition and a q-value of ≤0.05 were determined and the overlap between these gene lists displayed in Venn diagrams.

### TN-seq is a robust method to identify *S. aureus* genes involved in NaCl stress

In our experiments, NaCl stress elicited the strongest response. We therefore focused our further analysis on the *S. aureus* salt stress response and conducted two additional TN-seq screens to confirm the robustness of this approach (Figure 2A). TN-seq screens were performed using a previously published library (Santiago *et al*., 2015, Coe *et al*., 2019) with six transposon variants and the library containing only one transposon variant used in the first TN-seq experiment (Supplementary Table S1 for overview). The libraries were propagated for 17 generations in either LB or in medium containing 0.5 M NaCl (Fig. 2B). When the libraries were grown in high salt medium, clustering of transposon insertions in certain regions was observed due to the enrichment of better salt-adapted TN-strains (Supplementary Fig. S3). The fold-changes of transposon insertions per gene under the salt stress conditions compared to the LB condition (input library) as well as q-values were determined for each gene (Supplementary Table S6-S7). The data were plotted as volcano plots (Fig. 2C), revealing similar numbers of conditional salt essential and dispensable genes in both replicates. A good overlap of the top essential and dispensable genes was observed between the replicates, confirming the robustness of the TN-Seq screens (Supplementary Tables S8-11).

**Fig. 2.**
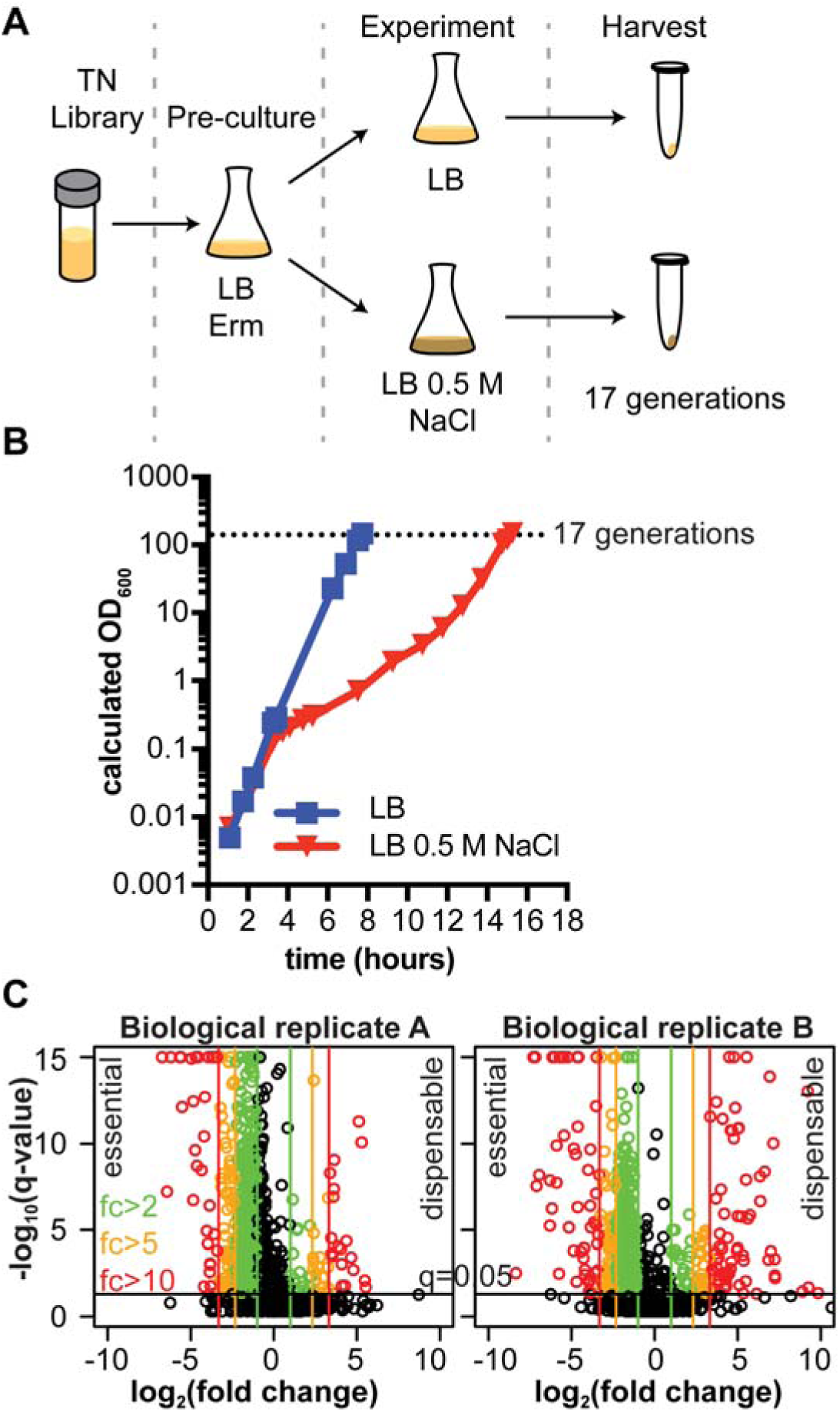
TN-Seq screen reveals essential and dispensable genes during prolonged NaCl stress in *S. aureus*. A. Workflow of TN-seq experiments. A saturated transposon library was pre-cultured for one hour, then used to inoculate LB medium containing either normal levels (0.086 M) or 0.5 M of NaCl. After 17 generations, cells were harvested, and transposon insertion sites determined by high throughput sequencing. B. Growth curve. Growth of the *S. aureus* library culture in LB and LB 0.5 M medium was followed by determining OD_600_ readings. Dotted line indicates the 17 generation threshold. Cultures were back-diluted once when they reached an OD of approximately 0.3 and the optical densities shown are calculated from measured ODs time dilution factor. Culture from replicate B (n=1) is shown and a similar growth profile was seen for replicate A. C. Volcano plots of essential and dispensable genes in NaCl conditions compared to LB. Negative log_2_ fold-changes indicate essential and positive log_2_ fold-changes dispensable genes. q-value stands for Benjamini-Hochberg false discovery rate. Black horizontal line indicates q-value of 0.05 (cut-off) and red, orange and green vertical lines 10-, 5- and 2-fold differences in either direction. Colored circles indicate genes for which a significant change in the number of transposon insertions was observed above q-value cutoff and follow the same color code as the vertical lines. Very small q-values were truncated to fit onto the graph. Data from replicate A are shown in the left panel and from replicate B in the right panel (n=1 for each plot).

### The TN-seq screens highlight the essentiality of transporters and cell envelope related genes and the dispensability of respiration genes under high salt conditions

In both replicates, genes coding for transporters and cell envelope related genes were the most prominent conditionally salt essential genes (Voronoi maps in Supplementary Fig. S4-S5, Top essential and dispensable genes in Supplementary Tables S8-S11). More specifically, the TN-seq data indicated that inactivation of the putative magnesium transporter MgtE (Schuster *et al*., 2019) and the putative D-serine/D-alanine/glycine transporter AapA is detrimental under salt stress (Voronoi maps in Supplementary Fig. S4, Top essential genes in Supplementary Tables S8 and S10). In addition, the *dlt* operon, coding for enzymes required for the D-alanylation of teichoic acids, an important factor for cationic peptide resistance (Brown *et al*., 2013), as well as the stationary and stress sigma factor gene sigB were found to be essential.

An even larger number of genes was identified as dispensable during salt stress (Voronoi maps in Supplementary Fig. S5, Top dispensable genes in Supplementary Tables S9 and S11). Inactivating or reducing the expression of such genes is expected to help *S. aureus* survive under salt stress. Among these dispensable genes were genes coding for the sodium antiporter Mnh2, which has previously been shown to transport Na^+^/H^+^ and K^+^/H^+^ in high pH medium and to play an important role in cytoplasmic pH maintenance (Vaish *et al*., 2018). As noted earlier, a large number of genes involved in respiration, such as the *cta/qox/men/hem* genes as well as genes involved indirectly in respiration through quinone synthesis (shikimate pathway, *aro operon*) were also identified as dispensable during salt stress. The data also reproducibly indicated that genes coding for the phosphodiesterases GdpP and Pde2 involved in degradation of the signaling molecule c-di-AMP are dispensable under high salt conditions. Using the TN-seq approach, we were able to identify a large number of *S. aureus* genes as potentially dispensable or essential when this organism is exposed to high salt.

### Tn-seq reveals previously unidentified salt resistance genes during long-term salt exposure

The TN-seq method relies on the enrichment of strains adapted to certain growth conditions and is therefore ideal for exploring the long-term effects on growth in response to salt stress. In a previous study (Price-Whelan *et al*., 2013), the adaptation of *S. aureus* to 2 M NaCl stress after 6 generations of growth was determined on a transcriptome level. This work highlighted the importance of K^+^ transporters for *S. aureus* to survive a high salt exposure. To evaluate how our results compare, we determined the overlaps between our datasets and the microarray data. To this end, the *S. aureus* strain COL locus tags used in the microarray study were converted to TCH1516 locus tags where possible (Supplementary Table S12) and compared to the two TN-seq datasets. The overlap between the downregulated (microarray) and dispensable (TN-seq) genes was minimal in both replicates (1 and 7) and also very small for the upregulated and essential genes (both 14). Three operons were consistently essential or upregulated between all datasets: the *cap5, SAUSA300_0771-2*, and the *sda* operon. The *cap5* genes are involved in capsule production (Chan *et al*., 2014) and have been previously reported to be induced by the addition of salt (Pöhlmann-Dietze *et al*., 2000). Genes *SAUSA300_0771-2* encode for membrane proteins that are predicted to form a threonine/serine exporter (Interpro database IDs: IPR010619 and IPR024528) and the *sda* genes are annotated as L-serine dehydratase components and a regulator protein. The low overlap between the microarray and TN-seq data (Supplementary Table S12) suggests that in our screen we have identified a number of previously unknown salt tolerance genes.

### Confirmation of genes essential during salt stress using defined mutants

To determine to what extent different genes identified in the TN-seq screen contribute to the salt resistance of *S. aureus*, a set of potentially salt essential genes from dataset B were chosen for further investigation (Supplementary Table S13). For this analysis we used defined transposon mutants available from the *S. aureus* NTML transposon mutant library (Fey *et al*., 2013). The growth of the different NTML transposon mutant strains was assessed in LB and LB 0.5 M NaCl medium. In LB medium, most strains grew similarly to the WT strain JE2, indicating that inactivation of the respective gene does not greatly affect bacterial growth in normal LB medium (Supplementary Figs. S6A-D, left panels). Exceptions were strains NE1109 (sigB), NE1778 (*lcpB*) and NE188 (mfd), cultures of which reached slightly lower final optical densities (Supplementary Figs. S6A-D, left panels). At high osmolality conditions (LB 0.5 M NaCl), all strains showed a growth defect compared to the WT strain (Supplementary Figs. S6A-D, right panels), validating the TN-seq as a method to identify *S. aureus* genes that are important during salt stress. Several strains exhibited strong growth defects in the presence of 0.5 M NaCl with strain NE1384, containing a transposon insertion in *SAUSA300_0957* showing extremely reduced growth (Supplementary Figs. S6C and D). To confirm that the salt sensitivity was mediated by the inactivation of the genes in questions, we constructed complementation plasmids for seven of the most promising candidates by either expressing the gene of interest from its native promoter or an anhydrotetracycline (Atet) inducible promoter in the transposon mutant strains. While no complementation was observed for four strains (Supplementary Figs. S7A and S7B), the salt-dependent growth defect could be complemented for strains carrying mutations in *SAUSA300_0694* (Fig. 3A), *SAUSA300_0910* (mgtE) and *SAUSA300_0957* (Fig. 3B). *SAUSA300_0694* encodes a hypothetical protein with 6 predicted transmembrane helices but no other identifiable domain motif. MgtE (SAUSA300_0910) is a predicted magnesium transporter, which we have recently shown can also contribute to cobalt toxicity in *S. aureus* (Schuster *et al*., 2019). *SAUSA300_0957* (from here on out referred to as gene *957*) codes for a cytoplasmic protein with a DUF2538 domain of unknown function that is conserved in Actinobacteria and Firmicutes. We further investigated this gene in this study because the *957* mutant exhibited the strongest salt-sensitivity phenotype (Fig. 3B).

**Fig. 3.**
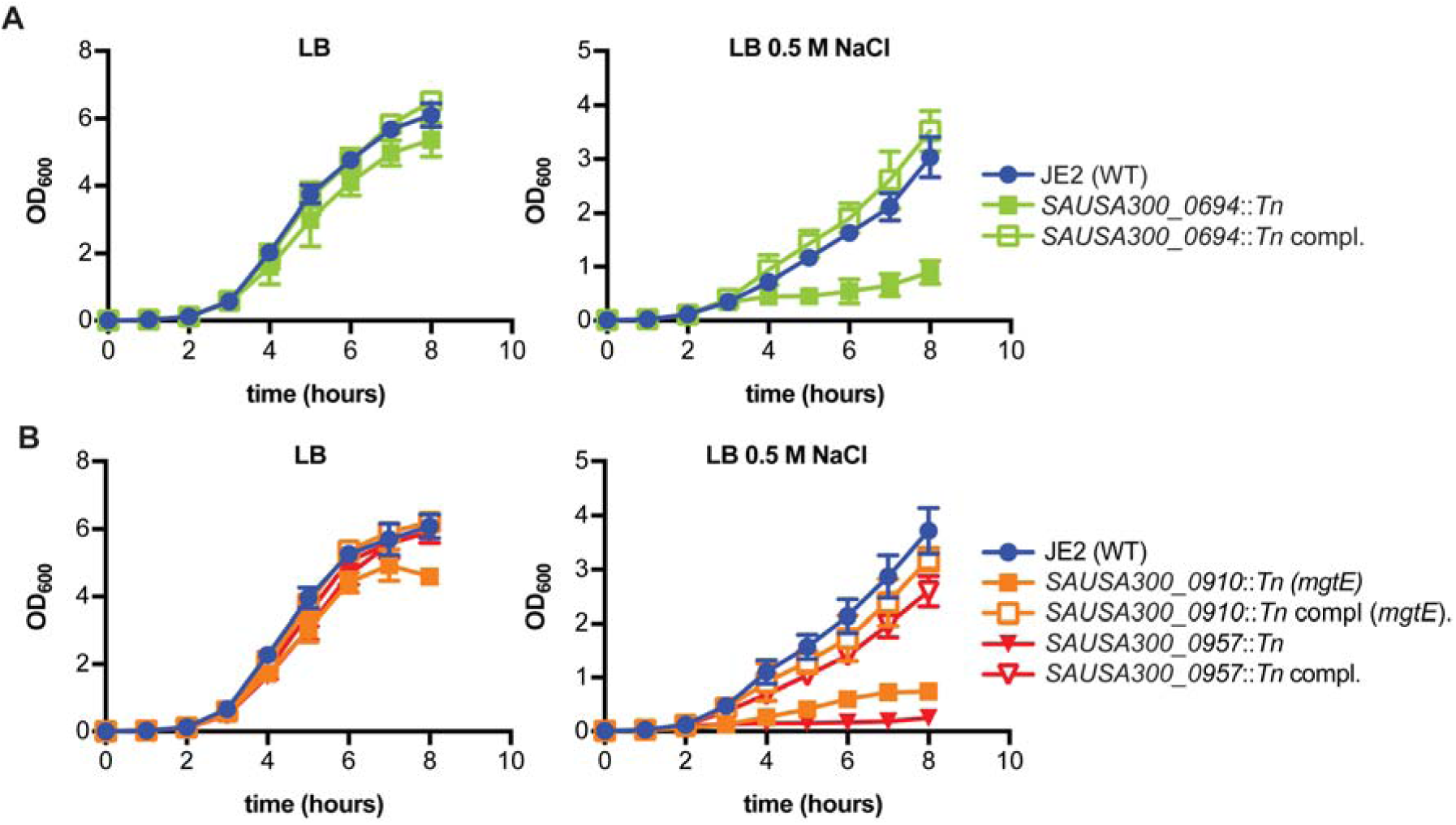
Growth curves and complementation analysis of *S. aureus* strains with transposon insertions in salt essential genes. A-B. *S. aureus* strains JE2 (WT), *SAUSA300_0694::Tn* mutant with empty plasmid and complementation strain (A) or JE2 (WT), *SAUSA300_0910 (mgtE), SAUSA300_0957::Tn* with empty plasmid (B) and respective complementation strains were grown in LB (left panels) or LB 0.5 M NaCl medium (right panels) supplemented with 100 ng/ml Atet and OD_600_ readings determined at timed intervals. Growth curves were performed in triplicates (n=3) and means and SDs of OD_600_ readings were plotted.

### Deletion of gene *957* causes physiological changes leading to a smaller cell size

As all previous experiments were performed with transposon mutants, we first constructed strain LAC*Δ*957* with a marker-less in-frame deletion of gene *957*. Strain LAC*Δ*957* had the expected growth defect in the presence of 0.5 M NaCl and this phenotype could be complemented by expressing a functional copy of *957* (Fig. 4A). To gain further insight into the molecular function of gene *957*, we analyzed the neighboring genomic area. Transcriptome data (Mäder *et al*., 2016) suggested that gene *957* is in an operon with *lcpB* (Fig. 4B), which is one of three wall teichoic acid (WTA) ligases present in *S. aureus* (Chan *et al*., 2013, Over *et al*., 2011, Schaefer *et al*., 2017). *fmtA* is located upstream of *957* and has been proposed to code for an esterase that can remove D-alanine modifications from teichoic acids (Rahman *et al*., 2016) and is involved in methicillin resistance (Berger-Bächi *et al*., 1989). Genes coding for a predicted acetyltransferase and *atl* coding for the major *S. aureus* autolysin are found immediately downstream of the *957* operon (Oshida *et al*., 1995). Due to the presence of many cell envelope related genes in the vicinity of gene *957*, we proceeded to test if the *957* mutant displays other phenotypes connected to cell envelope homeostasis. First, WT LAC*, the *957* mutant and the complementation strain were grown in LB and the ratio of colony forming units to optical density (CFU/OD) was determined. The CFU/OD ratio was significantly larger for the *957* mutant (60.8±6.8 ×10^7^ CFU/OD) mutant as compared to strain LAC* (37.5±9.3 ×10^7^ CFU/OD) when grown in LB, suggesting a change in cell shape or size (Fig. 4C). This phenotype was restored in the complementation strain (37.3±3.6 ×10^7^ CFU/OD) (Fig. 4C). When the average cell diameter was determined by microscopy, a small but significant reduction in cell diameter was seen for the *957* mutant (1.25±0.02 μm) as compared to the WT (1.36±0.03 μm) and the cell size was restored to wild type levels in the complementation strain (1.35±0.05 nm) (Fig. 4D). These results support our hypothesis that the envelope might be a target of *gene 957* and prompted us to look into these processes in more detail.

**Fig. 4.**
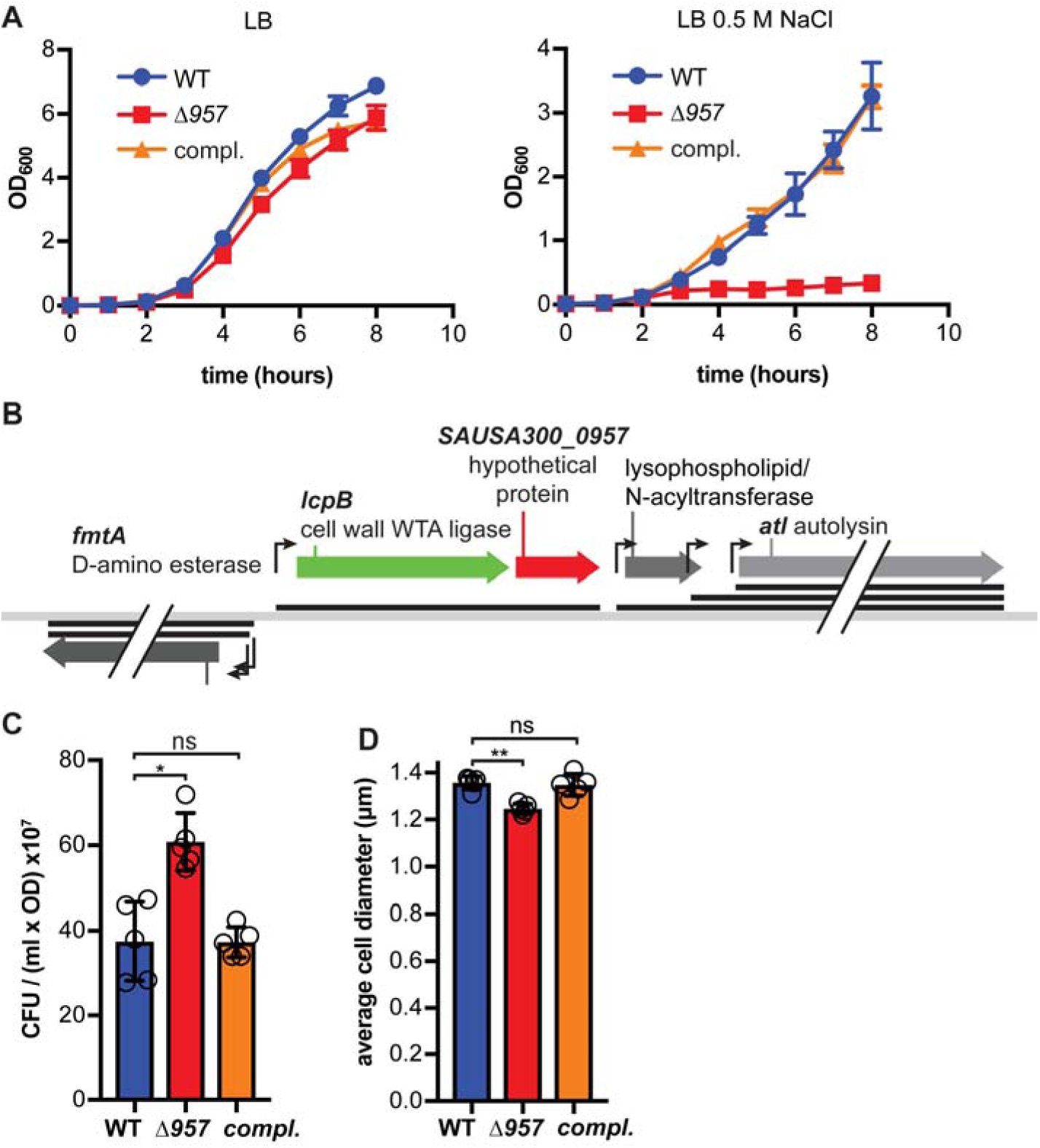
Gene *SAUSA300_0957* is located in a genomic region with cell envelope genes and its deletion leads to salt sensitivity and other phenotypic changes. A. Growth curves. *S. aureus* strains LAC* piTET (WT), LAC*Δ*957* piTET (Δ*957*) and the complementation strain LAC*Δ*957* piTET-*957* (compl.) were grown in LB or LB containing 0.5 M NaCl medium and 100 ng/ml Atet. The growth was monitored by determining OD_600_ readings the means and SDs from three independent experiments were plotted. B. Schematic of the *S. aureus* USA300 FPR3757 chromosomal region with gene *SAUSA300_0957* (gene *957*). Putative promoters are shown as angled arrows (adapted from (Mäder et al., 2016)) and black bars indicate the corresponding transcripts. C. Determination of CFU/OD ratios. OD_600_ values as well as CFU/ml were determined for overnight cultures of the *S. aureus* strains described in panel B and the means and SDs of the CFU per ml OD_600_ of 1 from five independent experiments were plotted. D. Cell size measurements. The cell walls of the *S. aureus* strains described in panel B were stained with fluorescently labelled vancomycin and the cells subsequently observed under a microscope. The diameters of 200 cells were measured and the means calculated. This average of the means and SDs from five independent experiments were plotted. For statistical analysis, a Kruskal-Wallis one-way ANOVA test was performed followed by a Dunn’s post-hoc test to determine p-values. Asterisks (*) indicate p≤0.05 and two asterisks (**) p≤0.01 and ns=not significant.

### Salt sensitivity of *957* is not due to interaction with LcpB but likely due to changes in peptidoglycan crosslinking

Since the gene *957* is on the same transcript as *lcpB* coding for a WTA ligase (Fig. 4B), the contribution of WTA to salt stress resistance as well as the contribution of *957* to the attachment of WTA to the cell wall was assessed. Initially the growth of a *tagO* (*tarO/llm*) mutant, a strain unable to produce WTA, was tested in LB and LB 0.5 M NaCl medium. When cells were grown in LB, they grew normally (Supplementary Fig. S8A), but under high salt conditions, the *tagO* mutant showed reduced growth, similar to that of the *957* mutant (Fig. 5A). This is consistent with what has been reported previously for a *Staphylococcus epidermidis tagO* mutant (Holland *et al*., 2011) and indicates an important role for WTA during salt stress. When the *lcpB* mutant was propagated in LB medium, its growth was slightly inhibited (Supplementary Fig. S8A). However, in high salt medium (Fig. 5A) it grew unimpeded and like WT. Since the *lcpB* mutant did not show a growth defect in high salt medium, this made it unlikely that *957* is a regulator of LcpB activity, and hence is involved in the attachment of WTA to the peptidoglycan. To specifically investigate the requirement of *957* in WTA attachment, we isolated WTA from strains grown in either LB or LB containing 0.4 M NaCl and separated the polymer on native polyacrylamide gels. The slightly lower concentration of salt was chosen because growth of the *957* and the *tagO* mutants was too strongly inhibited at 0.5 M NaCl. As expected the *tagO* mutant did not produce any WTA (Supplementary Fig. S8B, Fig. 5B) and consistent with previous studies (Over *et al*., 2011, Chan *et al*., 2013), the *lcpB* mutant exhibited slightly reduced WTA levels compared to the other strains (Supplementary Fig. S8B, Fig. 5B). No reduction in WTA was observable in the *957* mutant compared to the WT strain in either LB (Supplementary Fig. S8B & S8C) or LB 0.4 M NaCl (Fig. 5B & 5C) but rather a small increase, which could be complemented. Taken together, these data indicate that *957* neither directly nor indirectly (e.g. through regulating the activity of LcpB) plays a major role in WTA attachment.

**Fig. 5.**
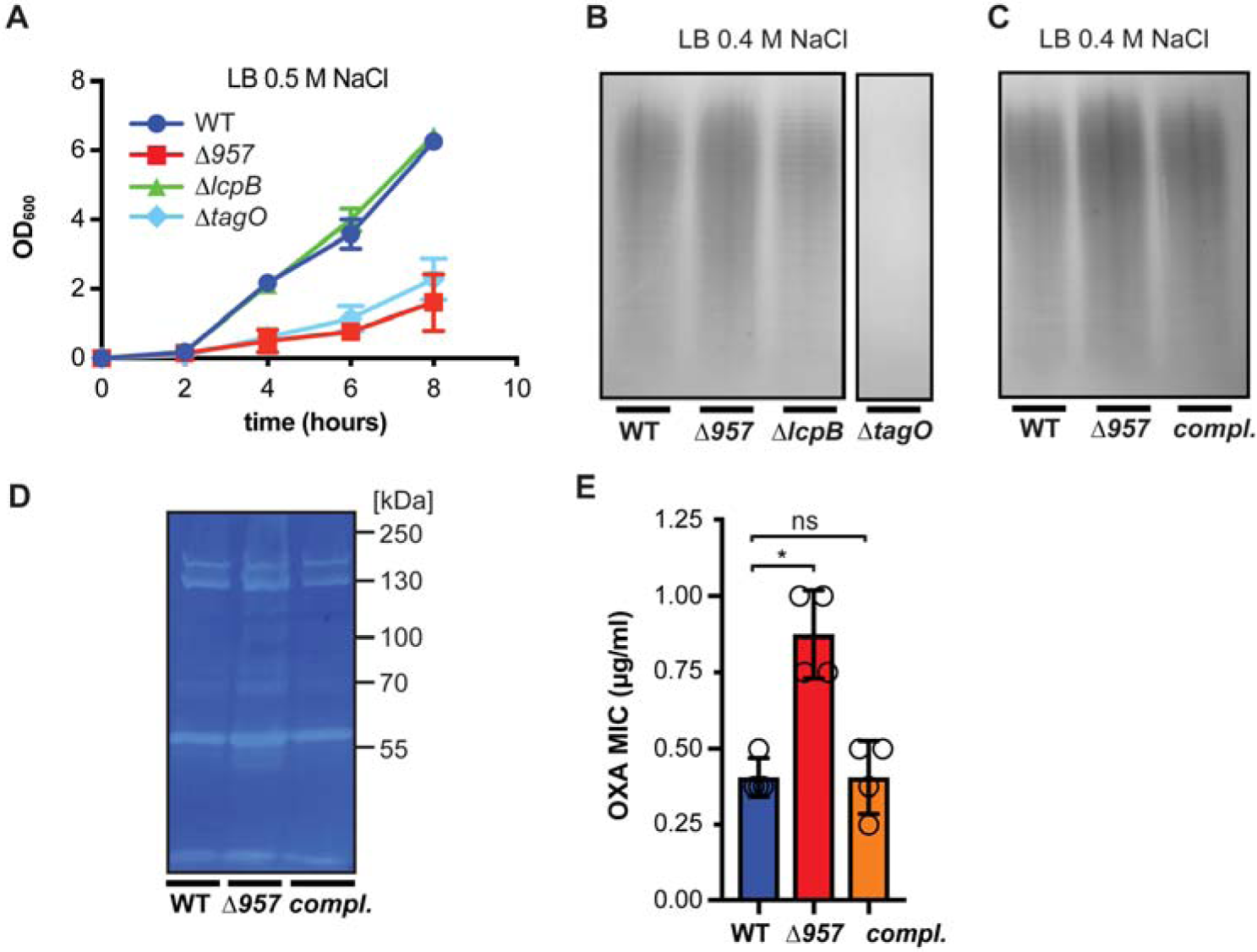
Inactivation of *gene 957* does not significantly alter WTA levels or autolysis but leads to changes in oxacillin resistance. A. Growth curves. *S. aureus* strains LAC* (WT), LAC*Δ*957* (Δ*957*), LAC*Δ*lcpB*, (Δ*957*), and LAC*Δ*tagO* (Δ*tagO*) were grown in LB 0.5 M NaCl medium and OD_600_ readings determined at timed intervals. The means and SDs from three biological replicates were plotted. B. Detection of WTA on silver stained gels. WTA was isolated from *S. aureus* strains described in panel A following growth in LB 0.4 M NaCl medium. The WTA was separated by electrophoresis and visualized by silver stain. The experiment was performed three times and one representative gel image is shown. C. Detection of WTA on silver stained gels. Same as panel B but using the strains LAC* piTET (WT), LAC*Δ*957* piTET (Δ*957*), and the complementation strain LAC*Δ*957* piTET-*957* (compl.) grown in LB 0.4 M NaCl medium also containing 100 ng/ml Atet. D. Zymogram gel. Cell extracts prepared from *S. aureus* strains described in panel C were separated on a gel containing heat killed *Micrococcus luteus* cells. Autolysins were renatured and the zones of lysis visualized by methylene blue staining. The experiment was performed twice and one experiment is shown. E. Determination of oxacillin MICs. Oxacillin MICs were determined for *S. aureus* strains described in panel C using Etest strips. The median and SDs from four biological replicates were plotted. For statistical analysis, a Kruskal-Wallis one-way ANOVA test was performed followed by a Dunn’s post-hoc test to determine p-values. Asterisks (*) indicate p≤0.05 and ns=not significant.

Gene *957* is found upstream of the bi-functional autolysin gene *atl* and therefore we next investigated potential changes in autolytic activity using zymograms. A small increase in autolytic activity was detected in the *957* mutant (Fig. 5D) for bands around 55 kDa and 70 kDa compared to the WT and complementation strain, indicating a possible change in Atl availability. We also tested the susceptibility of the *957* mutant towards the cell wall active beta-lactam antibiotic oxacillin. The *957* mutant exhibited a small but significant increase in MIC towards this antibiotic compared to the WT and the complementation strain (Fig. 5E), indicating potential changes to the peptidoglycan structure. To test this, we next analyzed the muropeptide profile of mutanolysin-digested peptidoglycan isolated from the WT, the *957* mutant and complementation strain after growth in LB or LB 0.4 M NaCl medium. Upon first inspection, no peaks were absent in any condition or strain (Supplementary Fig. S8D and S8E). Probing further, mono- and multimers of muropeptides up to 7-mers were quantified but no significant changes were found when strains were cultured in LB medium (Fig. 6A). At the 0.4 M NaCl condition however, the muropeptide profiles of the *957* mutant strain exhibited a significant decrease in the total amount of di- and trimers and a significant increase in higher multimers compared to the WT and complementation strain (Fig. 6B). This underlines the involvement of gene *957* in the cell wall homeostasis and suggests that the sensitivity to NaCl could potentially be caused by higher crosslinking and possibly rigidity of the peptidoglycan in the mutant.

**Fig. 6.**
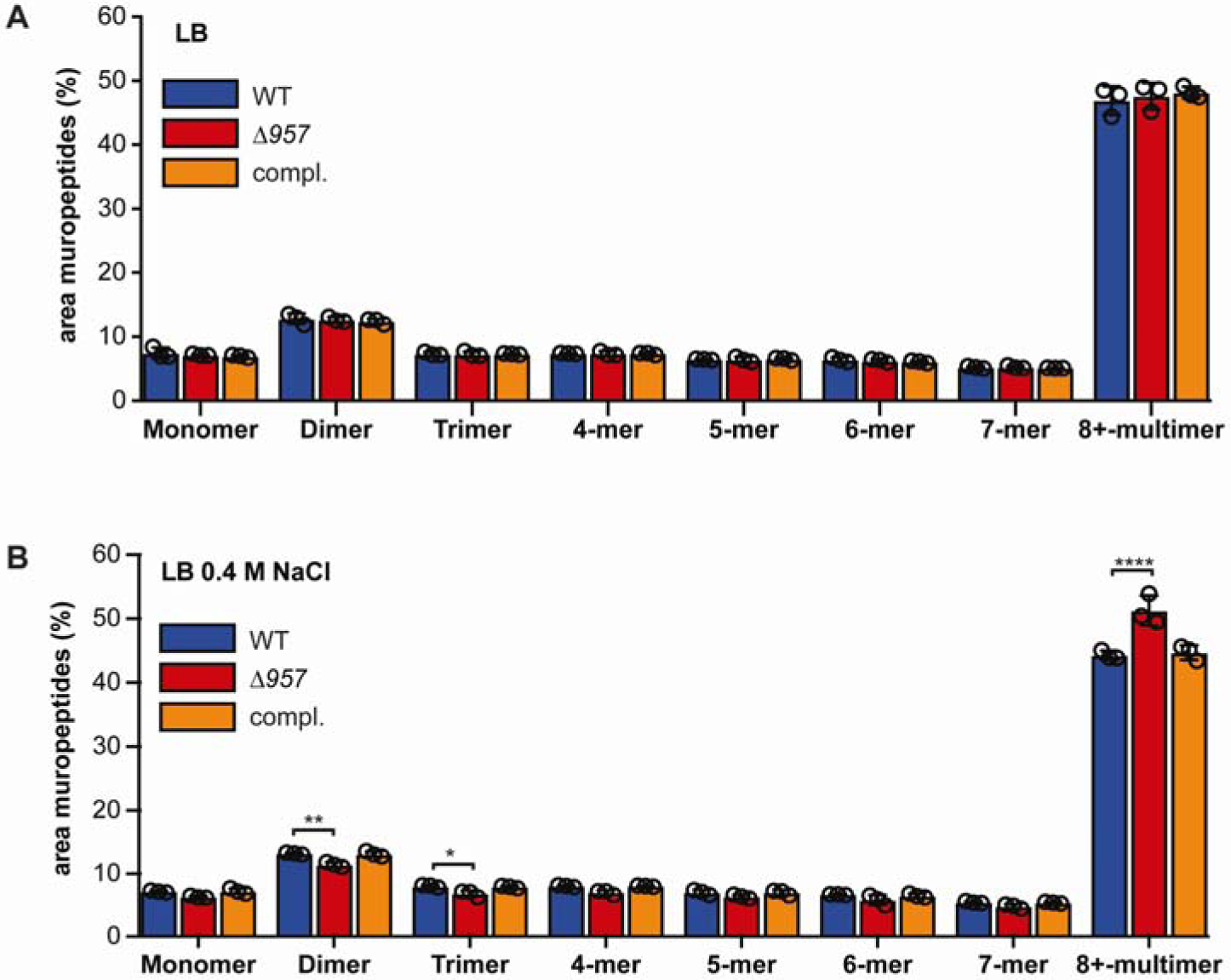
Inactivation of *gene 957* leads to an increase in peptidoglycan crosslinking. A-B. Muropeptide quantification. *S. aureus* strains LAC* piTET (WT), LAC*Δ*957* piTET (Δ*957*), and the complementation strain LAC*Δ*957* piTET-957 (compl.) were grown in either LB (A) or LB 0.4 M NaCl (B) containing 100 ng/ml Atet. After reaching late-exponential phase, the peptidoglycan was isolated, digested and the muropeptides separated by HPLC. The areas under the peaks from the chromatograms (representative chromatograms are shown in Supplementary Fig. S8) were quantified for each mono-, di- or multimer. Bars represent means, and error bars standard deviations from three independent experiments. For statistical analyses, two-way ANOVAs were performed, followed by Dunnett’s post-hoc tests using the WT as a comparison. The differences in (A) were not significant. For (B), one asterisk (*) indicates p≤0.05, two asterisks (**) p≤0.01 and four asterisks (****) p≤0.0001.

### Salt-resistant suppressors possess variations in the transglycosylase domain of Pbp2

Next, we attempted to generate *957* suppressor strains that showed improved growth in the presence of 0.5 M NaCl, with the idea that by mapping and investigating the compensatory mutations, further insight into the cellular functions of protein 957 could be gained. We readily obtained suppressor strains by growth of the *957* mutant strain in NaCl containing liquid medium. These strains grew as well as or even better than the WT LAC* strain in high salt conditions (Fig. 7A) and also grew similar to the WT in LB (Supplementary Fig. S9A). The genome sequences of 10 independent suppressor strains were determined and single nucleotide polymorphisms (SNPs) in one or two genes could be identified for each strain (Supplementary Table S14), except for one strain where the coverage was insufficient, and this strain was omitted from further analysis. A common denominator for all suppressor strains were SNPs in the *pbp2* gene, coding for the Penicillin binding protein 2 (Pbp2) (Fig. 7B). *S. aureus* Pbp2 is a bifunctional enzyme (Wyke, 1984, Murakami *et al*., 1994), which possesses transglycosylase and transpeptidase activity and is involved in peptidoglycan synthesis. All nine discovered SNPs were unique and led to mutations in the amino acid sequence in the transglycosylase domain but neither to stop codons, nor frameshifts nor mutations in the transpeptidase domain. The mutations were mapped (Fig. 7B) onto an available Pbp2 structure (3DWK) (Lovering *et al*., 2008) to see if a specific area of the transglycosylase domain had been targeted, but the amino acid substitutions were found throughout the molecule and not only in the active site. Interestingly, also from the Tn-Seq data, *pbp2* appeared to be considerably less important for growth at 0.5 M NaCl (Supplementary Fig. S9B and S9C), although transposon insertions were mainly located at the beginning and end of the gene or within the promoter region.

**Fig. 7.**
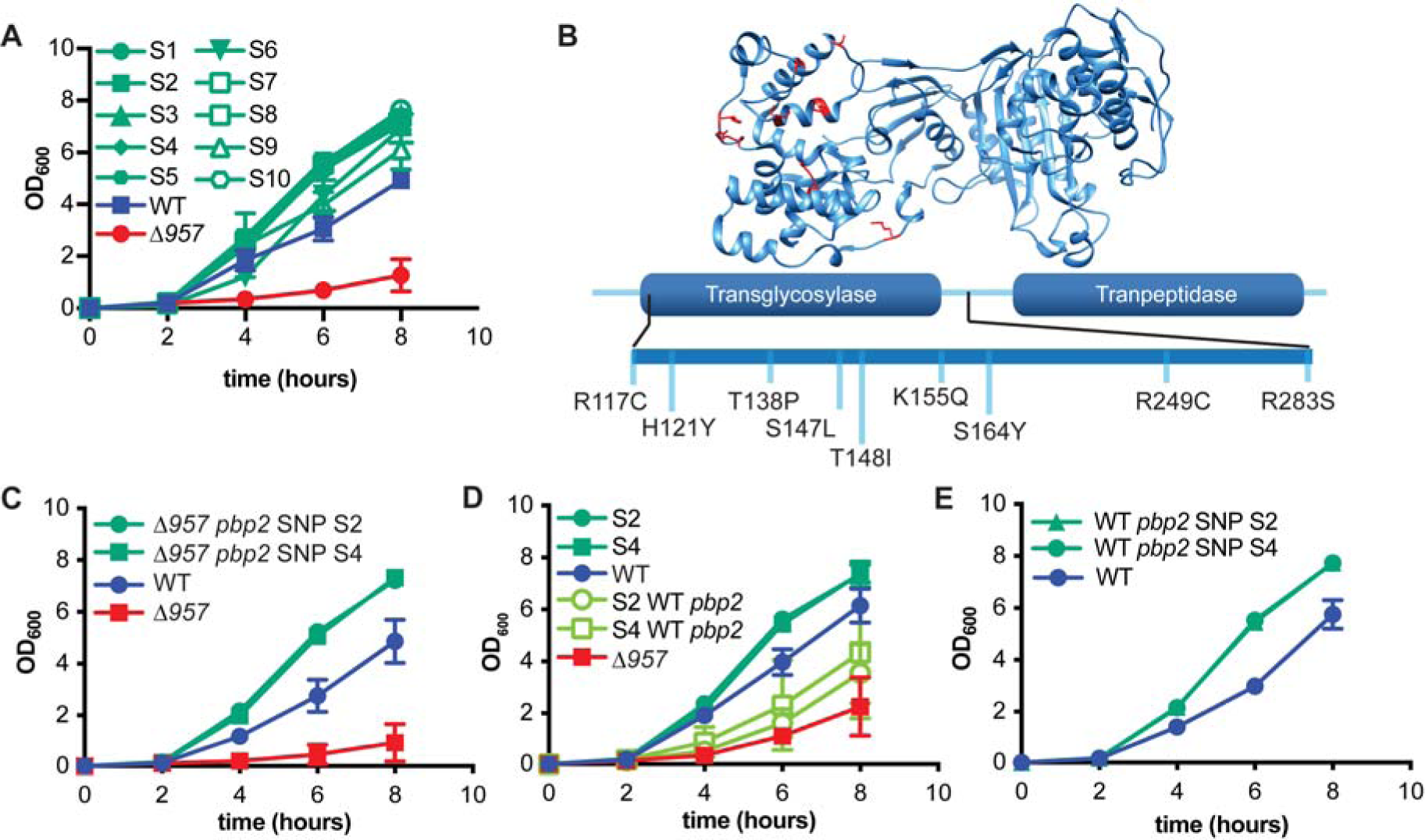
The growth and peptidoglycan defect observed for the *957* mutant can be rescued by compensatory mutations in *pbp2*. A. Growth curves. *S. aureus* strains LAC* (WT), LAC*Δ*957* (Δ*957*) and the LAC*Δ*957* suppressors S1-10 (S1 to 10) were grown in LB 0.5 M NaCl medium, OD_600_ readings determined and the means and SDs of four biological replicates plotted. B. Schematic of Pbp2 with amino acid substitutions identified. Top: Structure of the *S. aureus* penicillin binding protein Pbp2 (PDB: 3DWK, (Lovering *et al.*, 2008)) with amino acids that are altered in the obtained suppressor strains shown in red. Bottom: Schematic of the Pbp2 enzyme with the transglycosylase and transpeptidase domains as well as the observed amino acid changes in the transglycosylase domain indicated. C. Growth curves using *957* mutant strains containing *pbp2* alleles from suppressors S2 and S4. Growth curves were performed in LB 0.5 M NaCl medium and plotted as described in panel A but using *S. aureus* strains LAC* *1332*::Tn (WT), LAC*Δ*957 1332*::Tn (Δ*957*), LAC*Δ*957 1332*::Tn *pbp2* SNP S2 (Δ*957 pbp2* SNP S2), LAC*Δ*957* suppressor S4 *1332*::Tn (Δ*957 pbp2* SNP S4). D. Growth curves using *957* suppressors with their *pbp2* gene repaired to WT. Growth curves were performed in LB 0.5 M NaCl medium and plotted as described in panel A but using *S. aureus* strains LAC* *1332*::Tn (WT), LAC*Δ*957 1332*::Tn (Δ*957*), LAC*Δ*957* suppressor S2 *1332*::Tn (S2), LAC*Δ*957* suppressor *S4 1332*::Tn (S4), LAC*Δ*957* suppressor S2 *1332*::Tn repaired WT *pbp2* (S2 WT *pbp2*), and LAC*Δ*957* suppressor S4 *1332*::Tn repaired WT *pbp2* (S4 WT *pbp2*). E. Growth curves using WT strains carrying *pbp2* SNPs. *S. aureus* strains LAC* *1332*::Tn (WT), LAC* *1332*::Tn *pbp2* SNP S2 (WT *pbp2* SNP S2) and LAC* *1332*::Tn *pbp2* SNP S4 (WT *pbp2* SNP S4) were grown in LB 0.5 M NaCl medium and OD_600_ readings determined and the means and SDs of three biological replicates plotted.

To confirm that the *pbp2* SNPs were indeed responsible for the suppression of the salt sensitivity of the *957* mutant, we transferred the *pbp2* SNPs from two strains into a fresh LAC*Δ*957* background by co-transduction of a transposon in a nearby gene (Supplementary Fig. S9D, schematic). As expected, these “recreated” suppressor strains showed also increased salt resistance (Fig. 7C, Supplementary Fig. S10A). In addition, we repaired the *pbp2* gene in two suppressors to the WT *pbp2* allele and this reduced their ability to cope with NaCl stress, all consistent with the hypothesis that the SNPs in *pbp2* are responsible for the suppression phenotype (Fig. 7D, Supplementary Fig. S10B). When the same *pbp2* SNPs were transferred into a WT LAC* strain, a reproducible growth improvement of the SNP-bearing strains was observed in high salt medium, an indication that the *pbp2* mutations lead to a general growth improvement of *S. aureus* under NaCl stress conditions (Fig. 7E, Supplementary Fig. S10C). Overall these results show that several independent mutations in the *pbp2* gene can suppress the salt sensitivity of the *957* mutant and can improve growth of the WT strain.

### Mutations in the *pbp2* gene alter moenomycin susceptibility and result in reduced peptidoglycan crosslinking

Moenomycin is a phosphoglycolipid antibiotic that inhibits the transglycosylase activity of the *S. aureus* Pbp2 enzyme (reviewed in (Ostash & Walker, 2010)). To test if a decrease in Pbp2 transglycosylase activity results in improved growth of the *957* mutant in high salt conditions, cells were grown on LB agar containing 0.5 M NaCl with or without moenomycin. The growth behaviors of the WT strain and the *957* mutant were similar when grown on solid LB agar containing 0.5 M NaCl but lacking moenomycin (Fig. 8A, top panel). This discrepancy in the growth behavior of the *957* mutant in high salt liquid versus solid medium was already noted when we attempted to generate suppressors on agar plates. On the other hand, in medium supplemented with 0.02 μg/ml moenomycin, we observed better growth of the *957* mutant compared to the WT strain (Fig. 8A, lower panel). In addition, the *957* mutant also showed improved growth compared to the two suppressor strains S2 and S4 when moenomycin was added (Fig. 8A, lower panel). These findings are consistent with the idea that a partial inhibition of the glycosyltransferase activity of Pbp2 can improve the growth of the *957* mutant in high salt medium.

**Fig. 8.**
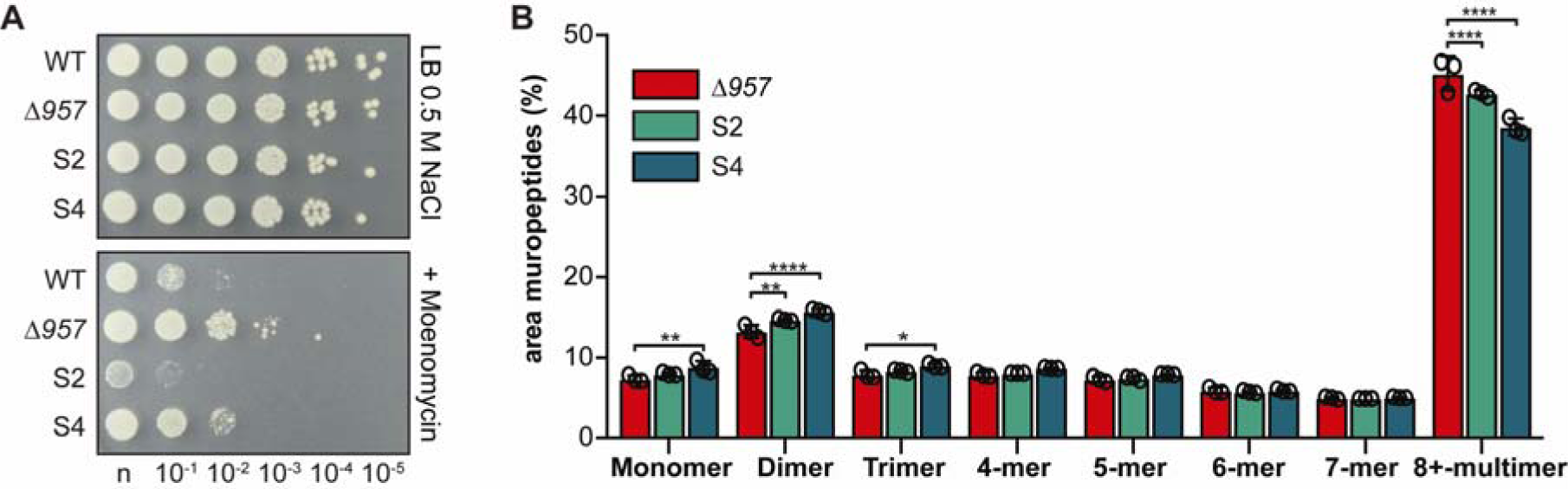
The Δ*957* suppressor strains show decreased peptidoglycan crosslinking compared the Δ*957* mutant. A. Bacterial growth on moenomycin supplemented agar plates. *S. aureus* strains LAC* (WT), LAC*Δ*957* (Δ*957*), LAC*Δ*957* Suppressor S2 (S2) and S4 LAC*Δ*957* Suppressor S4 (S4) were grown to exponential phase, normalized to an OD_600_ of 0.1 and plated either neat (n) or in 10-fold dilutions onto LB agar containing 0.5 M NaCl without (top) or with (bottom) moenomycin. Shown is one representative image of three biological replicates. B. Muropeptide analysis. *S. aureus* strains LAC*Δ*957* (Δ*957*), LAC*Δ*957* Suppressor S2 (S2) and S4 LAC*Δ*957* Suppressor S4 (S4) were grown in 0.4 M NaCl LB medium and muropeptides quantified as described in Fig. 5. Representative chromatograms are shown in Supplementary Fig. S10D. The means and SDs from three biological replicates were plotted. For statistical analysis a two-way ANOVA and Dunnett’s post-hoc test was performed. One asterisk (*) indicates p≤0.05, two asterisks (**) p≤0.01 and four asterisks (****) p≤0.0001.

Because the LAC*Δ*957* strain showed increased peptidoglycan crosslinking when compared to a WT strain, we hypothesized that the peptidoglycan of the suppressor strains would be again less crosslinked. To test this, the peptidoglycan from two of the suppressor strains grown in LB 0.4 M NaCl was isolated, the muropeptide profiles determined and compared to that of the original *957* mutant (Fig. 8B, Supplementary Fig. S10D). As before, the muropeptide profiles looked similar but quantification revealed a significant reduction in crosslinked peptidoglycan fragments and a significantly overrepresentation of monomeric and dimeric fragments in the suppressor strains as compared to the original *957* mutant (Fig. 8B). These results indicate that the amount of crosslinking of the peptidoglycan polymer is an important factor in the salt resistance of *S. aureus* and that dysregulation of peptidoglycan crosslinking can lead to destabilizing effects.

## Discussion

In this work, we performed TN-seq screens with *S. aureus* exposed to different osmotic stresses and could show that each osmotic stressor affects a defined but different set of genes. It is therefore important to use the term osmotic stress carefully, as our results highlight that there is not one general osmotic stress response but rather responses tailored to the individual osmolyte. Using the TN-seq method, we successfully identified several unknown salt tolerance genes in *S. aureus* and further characterized gene *957*, a gene without previously assigned function. Our results indicate that this gene product is involved in cell envelope homeostasis but likely not WTA attachment and we show that its absence leads to increased crosslinking of the peptidoglycan in the presence of NaCl. Suppressors of a *957* mutant acquired mutations in *pbp2* and the increase in peptidoglycan crosslinking was reversed in such stains. This demonstrates that the synthesis of the peptidoglycan layer is a tightly regulated process that plays an essential role during osmotic stress.

TN-seq experiments track the growth ability of single mutants in a mixed population and can therefore quickly and accurately determine genes that are required under certain growth condition. It is however always a concern that a single TN-seq experiment might not be reproducible due to stochastic extinction of individual mutants or certain growth dynamics. In this study, we therefore performed two independent NaCl TN-seq experiments using two different libraries and slightly different experimental setups. Although we detected differences between experiments, overall the data proved to be rather consistent suggesting that a single experiment would be sufficient in many cases.

From our TN-seq data and the use of different osmotic stressors (NaCl, KCl and sucrose), it is evident that although all these compounds exert similar osmotic stresses, the responses differs greatly between them (Figure 1 and Supplementary Figure S1-S2). NaCl and KCl conditions share a set of genes that become essential such as *aapA* and *mgtE* but differ in the set of dispensable genes. It is intriguing that a whole set of respiration related genes become dispensable under NaCl stress but not under KCl stress. This could indicate that respiration should be inhibited or slowed down in high NaCl conditions, possibly by interference with sodium pumps a process less likely to be inhibited by potassium ions. The set of essential genes under sucrose conditions was vastly different from that of the NaCl and KCl conditions whereas the set of sucrose dispensable genes looked similar to the KCl set. This highlights the differences in osmotic stress adaptations and serves as a reminder that the type of stressor remains important.

Most of the strains with mutations in genes we identified as NaCl essential in the TN-seq screens, proved to be sensitive to 0.5 M NaCl, confirming the effectiveness of the TN-seq screens (Supplementary Fig. S6). In particular, previously uncharacterized genes such as *SAUSA300_0694, mgtE*, and the *957* gene could be identified as essential during NaCl stress and their phenotypes could be complemented (Fig. 3). In our study, we opted not to experimentally validate any of the dispensable genes, as the verification of a growth advantage is more difficult to prove than a growth defect, but these genes will provide interesting starting points for future research. In addition, it will be interesting to investigate higher concentrations of NaCl to see if the same or different genes are flagged as essential and dispensable. Somewhat contradictory, our data indicated, that the c-di-AMP phosphodiesterase enzymes GdpP and Pde2 are dispensable when cells are exposed to high salt conditions. This is in contrast to the reports that construction and propagation of a *dacA* mutant (coding for the cognate c-di-AMP cyclase DacA) is only possible at high salt concentrations (or in defined chemical medium) (Zeden *et al*., 2018). These findings do however point to an involvement of c-di-AMP in salt mediated osmotic stress adaption in *S. aureus*.

Gene *957* was selected for further characterization as part of this study, because it had previously not been linked to NaCl stress and had not been investigated before. Initial tests supported our proposed link to the cell envelope. The differences in CFU/OD correlations of the WT and *957* mutant could be explained by the reduced cell size of the *957* mutant, as this would produce more cells per OD unit. The underlying mechanism as to why *957* mutant cells are smaller in size is still unclear but could be related to changes in peptidoglycan homeostasis.

As gene *957* is co-transcribed with *lcpB*, we initially assumed the encoded protein is involved in WTA attachment via LcpB. However, we were unable to demonstrate such a link. Instead the *957* mutant exhibited increased muropeptide crosslinking in the presence of 0.4 M NaCl, which could be suppressed by *pbp2* SNPs suggesting an involvement in peptidoglycan synthesis. We can only speculate if the SNPs identified in *pbp2* increase or inhibit the activity of Pbp2. However, judging by the number of independent SNPs, it seems more likely that the mutations lead to a decrease rather than an increase in transglycosylase activity. In addition, the results from the moenomycin sensitivity experiments using a sublethal antibiotic concentration highlights that partial inactivation of the transglycosylase activity of Pbp2 improves the growth of the *957* strain in high salt conditions. This is consistent with the idea that the obtained *pbp2* SNPs lead to a decrease in transglycosylase activity. The absence of frameshift or non-sense mutations can be explained by the importance of the C-terminal transpeptidase domain of *pbp2* (Pinho *et al*., 2001) and this is also reflected in the transposon insertion distribution in *pbp2*. Intriguingly, the suppressor mutants show reduced growth compared to the *957* mutant on moenomycin plates, presumably due to transglycosylation activity being reduced to levels that are detrimental to the cell. At this point it is unclear how mutations in the transglycosylase domain alter the muropeptide pattern, since the peptide bonds are made by the transpeptidase, not the transglycosylase domain. We hypothesize that a decrease in the efficiency of the glycosylation activity and slowing down the glycan chain synthesis process will reduce the efficiency of the subsequent transpeptidation process, resulting in decreased peptidoglycan crosslinking in the *pbp2* SNP strains. It is also noteworthy that the *pbp2* gene becomes dispensable under NaCl conditions, an additional indicator that Pbp2 activity needs to be avoided under high salt conditions.

Interestingly, the *957* mutant exhibited a slightly higher oxacillin MIC than the WT. This is in contrast to what we expected as MICs are determined in 2% (0.342 M) salt medium and this should lead to a growth disadvantage of the mutant. However, when seen in light of the increase in peptidoglycan crosslinking, this could explain the growth advantage of the *957* mutant in the presence of a beta-lactam antibiotic, as this could counter the inhibition of transpeptidases by oxacillin.

Although we were unable to determine the exact molecular mechanism, it is clear from our data that gene *957* is involved in cell wall synthesis either directly or indirectly. Based on these data, we hypothesize that the rigidity of the peptidoglycan cell wall could be an important factor to counter osmotic stresses, but further experimental work is needed to explore this in depth.

In conclusion, we have demonstrated differences between the osmotic stress response to NaCl, KCl and sucrose in *S. aureus*. In addition, we identified a number of previously uncharacterized factors involved in the osmotic stress response of *S. aureus*, which in particular highlighted the importance of the cell envelope. The generated data will also provide a great resource for further studies on the staphylococcal NaCl stress response.

## Experimental procedures

### Growth of bacteria

*E. coli* and *S. aureus* strains (see Supplementary Table S15 for details) were streaked from frozen stocks onto Lysogenic Broth (LB) or Tryptic Soya agar (TSA) plates, respectively. For all *E. coli* and most *S. aureus* experiments, the bacteria were grown in LB medium (Lennox Recipe: 10 g Tryptone, 5 g Yeast extract, 5 g NaCl) pH 7.5 or LB medium with extra 0.5 M NaCl, 0.5 M KCl or 1.0 M Sucrose added. For molecular cloning and some other experiments (as indicated in the text), *S. aureus* strains were grown in TSB medium. Growth curves were performed in 125 ml flasks with 20 ml of medium without antibiotics. Cultures were inoculated from overnight cultures to a starting OD_600_ of 0.01 and growth was followed by determining OD_600_ readings every one or two hours.

### TN-seq experiments

A previously published highly-saturated *S. aureus* USA300 library with a mix of six different transposons was used (Santiago *et al*., 2015, Coe *et al*., 2019). In addition, a new transposon library with only the blunt transposon, was produced using the same techniques as described (Santiago *et al*., 2015) containing approximately 1.2 million individual clones.

In the first experiment (NaCl, KCl and sucrose stress), a tube of the blunt library was thawed on ice and used to set up a 20 ml pre-culture in LB/Erm 10 μg/ml with a starting OD_600_ of 0.1. This starter culture was grown for 1 hour and subsequently used to inoculate 25 ml of either LB (Lennox Recipe: 10 g Tryptone, 5 g Yeast extract, 5 g NaCl) pH 7.5 or LB with extra 0.5 M NaCl, extra 0.5 M KCl or extra 1 M sucrose without Erm to a starting OD_600_ of 0.00125. When the cultures reached an OD_600_ of approximately 0.3, the cultures were back-diluted 1:100-fold into 40 ml fresh medium and 10-12 ml harvested by centrifugation when the back-diluted culture reached an OD_600_ of approx. 1.0 (16 generations). The bacterial pellets were washed once with their respective growth medium and stored at -20°C for further processing.

In the second experiment (NaCl, Replicate A), the multiplexed promoter transposon library was used and Erm 5 μg/ml was added to the pre-culture and subsequent cultures in either LB (Lennox Recipe: 10 g Tryptone, 5 g Yeast extract, 5 g NaCl) pH 7.5 or LB medium containing a final concentration of 0.5 M NaCl. Samples were collected at an OD_600_ of about 1.4 (17 generations). The third experiment (NaCl, replicate B) was set up similar to replicate A, however the blunt library was used in this case and Erm 5 μg/ml was only added to the pre-culture.

The library preparation for sequencing was done as described previously (Santiago *et al*., 2015). Briefly, genomic DNA was isolated, cut with *NotI*, biotinylated adapters ligated, the DNA cut again with *MmeI*, and fragments ligated to Illumina adapters and the products PCR amplified, incorporating bar codes and the Illumina adaptor sequences P5 and P7. The samples were sequenced on an Illumina HiSeq machine after spiking with 40% PhiX DNA. The data analysis was done using the Tufts Galaxy Server and custom python scripts as described earlier (Santiago *et al*., 2015). To this end, the reads were trimmed up to the barcode and de-multiplexed by strain barcode. Due to the variability in the DNA cleavage by the *MmeI* restriction enzyme, the reads that could not be mapped to a barcode were trimmed by an additional base and the process repeated. Reads were then mapped to the USA300_TCH1516 (CP000730.1) genome and a hop table was generated. Statistical analysis (Mann-Whitney tests and Benjamini-Hochberg) was done using custom python scripts (Santiago *et al*., 2015). Further exploration of the data was done using Excel and custom R scripts.

### *S. aureus* cell diameter measurements

For cell diameter measurements, *S. aureus* strains ANG4054, ANG4340, ANG4341 were grown overnight (14-20 hours) at 37°C with 180 rpm shaking in 5 ml LB in the presence of 100 ng/ml Atet. 100 μl of these overnight cultures was transferred to a 1.5 ml reaction tube and 1.5 μl of BODIPY-FL-vancomycin (100 μg/ml in phosphate buffered saline) was added to each tube to stain the cell wall. After a 30 min static incubation step at 37°C, 1.5 μl of the suspension was pipetted onto a slide covered with a 1% agarose pad and analyzed by microscopy using an Axio Imager.A2 microscope with an EC Plan-Neofluar 100x/1.30 Oil M27 objective and images recorded using an AxioCam MR R3. Native CZI files were opened in FIJI (Schindelin *et al*., 2012) and cell diameters of 200 cells were measured using the line and measure tool. Only cells without any visible septa were measured. Experiment was conducted five times with independent biological cultures (n=5, 200 cells each).

### CFU/OD correlations

The different *S. aureus* strains were grown as described for the cell diameter measurements. Optical densities of cultures were measured as well as 100 μl of 10^-6^ dilutions (made in their respective growth medium) plated onto TSA plates. The next day, the number of colonies were counted and the ratios of OD_600_ to CFU calculated. The experiment was conducted six time with independent biological samples (n=6).

### Zymogram assays

The different *S. aureus* strains were grown over night in TSB. The next day, the bacteria from an OD_600_ equivalent of 20 were pelleted by centrifugation for 3 min at 17000 × *g*. The cells were washed twice with 600 μl PBS and subsequently suspended in 50 μl SDS sample buffer. The suspension was boiled for 20 min with interspersed shaking, centrifuged for 5 min at 17000 × *g* and 20 μl of each sample was loaded onto Zymogram gels, which where were prepared as described previously (Atilano *et al*., 2014). Gels were stained and de-stained using methylene blue and water. The experiment was performed twice, and a representative result is shown.

### WTA isolation and analysis

Overnight cultures of *S. aureus* strains ANG4054, ANG4340, ANG4341, ANG1575, ANG4290, ANG4748, ANG4749, ANG4759 were prepared in LB and used to inoculate 50 ml LB or LB 0.4 M NaCl to an OD_600_ of 0.01. The cultures were incubated at 37°C with shaking until they reached an OD_600_ of 5 to 6. Where appropriate, 10 μg/ml chloramphenicol and 100 ng/ml Atet was also added to the medium. The bacteria from 20-24 ml culture were harvested by centrifugation for 10 min at 7000 *x g.* The bacterial pellets were washed with 50 mM 2-(N-morpholino) ethanesulfonic acid (MES) pH 6.5 buffer and stored at -20°C until further use. Pellets were then processed and WTA separated on a 20% native polyacrylamide gel by electrophoresis on a Biorad Protein XL ii cell as described previously (Covas *et al*., 2016) and WTA visualized by silver staining according to the manufacturer protocol. Experiments were done with three biological replicates (n=3) and a representative result is shown.

### Peptidoglycan isolation and analysis

Peptidoglycan was prepared as described previously (Corrigan *et al*., 2011) with the following modifications: 0.5 L (ANG4290, ANG4382, ANG4384) or 1.0 L (ANG5054, ANG4340, ANG4341) of LB 0.4 M NaCl medium (all strains) or LB (ANG5054, ANG4340, ANG4341) medium were inoculated to an OD_600_ of 0.01. For strains carrying derivatives of plasmid piTET, the medium was supplemented with 100 ng/ml Atet starting from the pre-cultures. The cultures were grown at 37°C with shaking at 180 rpm until they reached an OD_600_ of 2-3 and bacteria were subsequently harvested by centrifugation. Chromatography of mutanolysin digested peptidoglycan was performed as described previously using an Agilent 1260 infinity system (Corrigan *et al*., 2011) and muropeptide peaks assigned according to de Jonge *et al*. (de Jonge *et al*., 1992). For the muropeptide quantification, a baseline was drawn, the total peak area determined as well as the peak areas for mono- and the different multimer peaks and calculated as percentage of the total peak area. The peak area quantification was done three times for each HPLC chromatogram and average values were calculated. Experiments were done with three biological replicates (n=3).

### Data analysis, software and statistics

Data were processed with a combination of Python 3.6 (https://www.python.org), R 3.3 & 3.5 (https://www.r-project.org/), RStudio 1.1 & 1.2 (https://www.rstudio.com/), Prism 7 and 8 (https://www.graphpad.com), ChemStation OpenLab C.01.05 (https://www.agilent.com/) and Microsoft Excel 15 and 16 (https://www.office.com/). Microscopy images were analyzed using FIJI 1.0 (https://fiji.sc/). Statistical analysis was performed with Prism using appropriate tests as described in the figure legends.

### Raising of Δ*957* suppressor strains

Multiple independent overnight cultures of strain LAC*Δ*957* (ANG4290) were prepared in 5 ml LB. These cultures were back diluted the next day into 20 ml LB 0.5 M NaCl to a starting OD_600_ of 0.05 and grown for 8-10 hours at 37°C with shaking until the cultures were slightly turbid. 50 μl of each culture was passed into 5 ml of LB and grown overnight. The next day, appropriate culture dilutions were plated onto LB agar plates and incubated overnight at 37°C. For each independent culture, multiple single colonies were picked and used to inoculate individual wells of a 96-well microtiter plates containing 100 μl LB and the plates were subsequently incubated at 37°C in a 96-well plate incubator with shaking at 650 rpm. The next day, a culture aliquot was stored at -80°C, and the cultures were also diluted 1:50 in LB 0.5 M NaCl medium and 20 μl of these diluted cultures used to inoculate 180 μl of LB 0.5 M NaCl medium. The plates were incubated overnight with shaking at 37°C and the next morning the growth of the potential 957 suppressor strains compared to that of the original LAC*Δ*957* deletion strain (ANG4290), which showed low growth and the WT LAC* (ANG1575) strain, which showed good growth. For each lineage, four suppressors that showed good growth were streaked out from the previously frozen stocks and subsequently single colonies selected to set up overnight cultures. The deletion of gene *SAUSA300_0957* was confirmed by PCR and after performing growth curves in culture flasks, one strain that showed significant growth improvement compared to the original LAC*Δ*957* (ANG4290) strain was selected from each lineage. These strains were streaked out again for single colonies and used to inoculate an overnight culture, giving rise to the independently raised suppressor strains ANG4381 through ANG4394. The compensatory mutations for several of these suppressor strains were subsequently determined by whole genome sequencing.

### Transfer of *pbp2* SNPs by phage transduction

In order to demonstrate that the SNPs in *pbp2* are responsible for the growth rescue of strain LAC*Δ*957* (ANG4290), we transferred two different *pbp2* SNPs by phage transduction into the original LAC*Δ*957* strain as well as repaired these SNPs in the suppressor strains. This was done by placing an Erm-marked transposon in proximity of the *pbp2* gene allowing at a certain frequency for the co-transduction of the Erm marker and the *pbp2* SNP. To this end, the NTML transposon mutant library strain NE789 (Fey *et al*., 2013) was used, which harbors a transposon insertion in *SAUSA300_1332* (putative exonuclease) about 10 kbp upstream of the *pbp2* gene. *SAUSA300_1332* is expected to be unrelated to salt stress and the cell wall synthesis machinery but close enough to lead to an intermediate rate of co-transference with the *pbp2* SNPs. The transposon from NE789 was transduced into WT LAC* (Boles *et al*., 2010) and strain LAC*Δ*957* (ANG4290) using phage Φ85 yielding strains ANG4561 and ANG452, respectively. The transposon was also transduced into the suppressor strains ANG4382 and ANG4384 transferring either *SAUSA300_1332::Tn* only, yielding strains ANG4527 and 4528 or transferring *SAUSA300_1332::Tn* as well as replacing the *pbp2* SNP with a WT *pbp2* allele yielding *pbp2* repaired strains ANG4557 and ANG4568. Lysates from strains ANG4527 and 4528 (suppressors containing *pbp2* SNPs and *SAUSA300_1332::Tn*) were used to transfer the *pbp2* SNPs back into a clean LAC*Δ*957* strain background yielding strains ANG4624 and ANG4625 and into WT LAC*, yielding strains ANG4563 and ANG4564. Successful repair or transfer of the SNPs was checked by PCR and subsequent restriction digest of the product choosing enzymes that recognize sites present in either the WT or SNP allele (ANG4382 *pbp2* SNP: BsrI. Recognition site CCWGG, additional site introduced by SNP; ANG4384 *pbp2* SNP: SspI. Recognition site: AATATT, site missing in SNP).

### Determination of oxacillin MICs using Etest strips

*S. aureus* strains LAC* piTET (ANG4054), LAC*Δ*957* piTET (ANG4340) and the complementation strain LAC*Δ*957* piTET-*957* (ANG4341) were grown overnight (22 hours) in 5 ml TSB containing 10 μg/ml chloramphenicol and 100 ng/ml Atet. The next day, the cultures were diluted to an OD_600_ of 0.1 in sterile PBS buffer and streaked with cotton swabs onto cation adjusted Mueller-Hinton agar plates supplemented with 2% NaCl and 100 ng/ml Atet. One M.I.C.Evaluator strip was placed on each plate and the plates were incubated for 24 hours at 35°C. MICs were then read directly from the strips. The experiment was done with 4 biological replicates (n=4).

### Moenomycin growth improvement test

*S. aureus* strains LAC* (ANG1575), LAC* Δ*957* (ANG4290), LAC* Δ*957* S2 (ANG4382) and LAC* Δ*957* S4 (ANG4384) were grown overnight (18 hours) in 5 ml LB. The next day, the cells were diluted to an OD_600_ of 0.01 in LB, grown until mid-exponential phase (OD_600_ 0.4-0.6) and normalized to an OD_600_ of 0.1. The cells were 10-fold serially diluted and 5 μl of each dilution spotted onto LB agar containing 0.5 M NaCl with either 0.02 μg/ml moenomycin (mix of moenomycin A, A12, C1, C3 and C4, Santa Cruz Biotechnology) or no moenomycin. Plates were incubated at 37°C overnight (18-22 hours) and photographed. The experiment was done with 3 biological replicates (n=3) and one representative result is shown.

### Whole genome sequencing

Genomic DNA was either isolated using the Promega Gene Wizard kit according to the manufacturer instructions or using chloroform-isoamylalcohol as described previously (Schuster & Bertram, 2014). DNA was sent off for whole genome sequencing to MicrobesNG, Birmingham, U.K. or libraries prepared using the Illumina Nextera DNA kit and sequenced at the London Institute of Medical Sciences. Short reads were trimmed in CLC workbench genomics (Qiagen), then mapped to a custom *S. aureus* USA300 LAC* reference genome generated in a previous study (Bowman *et al*., 2016) and single nucleotide polymorphisms called based on at least 80% frequency of occurrence. This list was compared to a manually curated list of well non false-positives and these entries were removed.

### Nebraska Transposon Mutant Library (NTML) strains and complementation strains

All Nebraska transposon mutant library (NTML) strains and primers used in this study are listed in Supplementary Tables S13, S15 and S16. Transposon insertions in the respective genes were confirmed by PCRs. For complementation analysis, NTML strains NE535 (JE2 *SAUSA300_0867::Tn*), NE867 (JE2 *SAUSA300_0483::Tn*) were transformed as controls with the empty plasmid pCL55 (Lee *et al*., 1991), yielding strains ANG4326 and ANG4328 and strains NE188 (JE2 *SAUSA300_0481::Tn*), NE251 (JE2 *SAUSA300_0482::Tn*), NE526 (JE2 *SAUSA300_0694::Tn*), NE736 (JE2 *SAUSA300_0910::Tn*), NE1384 (JE2 *SAUSA300_0957::Tn*) with plasmid piTET (Gründling & Schneewind, 2007), yielding ANG4308, ANG4310, ANG4377, ANG4336, ANG4338. The WT strain JE2 (Fey *et al*., 2013) was also transformed with both plasmids yielding JE2 pCL55 (ANG4325) and JE2 piTET (ANG4307).

The complementation plasmid piTET-*481* for strain NE188 (JE2 *SAUSA300_0481::Tn*) was constructed by amplifying the *SAUSA300_0481* gene using primers P2378/P2379, digesting the product with AvrII/BglII and ligating the fragment with plasmid piTET cut with the same enzymes. The plasmid was then introduced into *E. coli* strain XL1-Blue, yielding ANG4139. After shuttling the plasmid through *E. coli* strain IM08B (Monk *et al*., 2012), creating strain ANG4150, the plasmid was integrated into the geh locus of NE188 (JE2 *SAUSA300_0481::Tn*) yielding strain ANG4309. The complementation plasmid piTET-*482* for strain NE251 (JE2 SAUSA300_0482::Tn), piTET-*694* for strain NE526 (JE2 *SAUSA300_0694::Tn*), piTET-*910* for strain NE736 (JE2 *SAUSA300_0910::Tn*), and piTET-*957* for strain NE1384 (JE2 *SAUSA300_0957::Tn*) were constructed in a similar manner using primer pairs P2380/P2381 (*SAUSA300_0482*), P2388/P2389 (*SAUSA300_0694*), P2384/P2385 (*SAUSA300_0910*) and P2386/P2387 (*SAUSA300_0957*), respectively. The complementation plasmids were recovered in *E. coli* XL1-Blue yielding strains ANG4140 (*SAUSA300_0482*), ANG4142 (*SAUSA300_0694*), ANG4144 (*SAUSA300_0910*) and ANG4145 (*SAUSA300_0957*), shuttled through *E. coli* IM08B giving strains ANG4151 (*SAUSA300_0482*), ANG4153 (*SAUSA300_0694*), ANG4155 (*SAUSA300_0910*) and ANG4156 (*SAUSA300_0957*) and finally introduced in the respective NTML strain yielding the complementation strains ANG4311 (*SAUSA300_0482*), ANG4378 (*SAUSA300_0694*), ANG4337 (*SAUSA300_0910*) and ANG4339 (*SAUSA300_0957*).

Complementation plasmid pCL55-*483* for strain NE867 (JE2 *SAUSA300_0483::Tn*) was made by fusing PCR products of the operon promoter (in front of *SAUSA300_0481*) amplified with primers P2588/P2553 and the *SAUSA300_0483* gene, amplified with primers P2554/P2589 together using primers P2588/P2589. The resulting fragment was cloned into pCL55 using EcoRI/BamHI restriction sites. The plasmid was recovered in XL1-Blue, creating strain ANG4291, shuttled through *E. coli* IM08B (ANG4292) and subsequently introduced into NE867 (JE2 *SAUSA300_0483::Tn*) to create strain ANG4327. The complementation plasmid pCL55-*867* for strain NE535 (JE2 *SAUSA300_0867::Tn*) was constructed in a similar manner, using primers P2590/P2556 and P2557/P2591 to amplify the promoter and *SAUSA300_0867* gene, which were subsequently fused in a second PCR using primers P2590/P2591. The resulting fragment was cloning into pCL55 using EcoRI/BamHI restriction sites and the resulting plasmid recovered in *E. coli* XL1-Blue (ANG4293), shuttling through IM08B (ANG4294) and finally introduced into NE535 (JE2 *SAUSA300_0867::Tn*), creating the complementation strain ANG4329.

### Construction of *S. aureus* gene deletion and complementation strains

*S. aureus* strains with in-frame gene deletions were constructed by allelic exchange using plasmids pIMAY (Monk *et al.*, 2012) and pIMAY* (Schuster *et al*., 2019). The gene deletion plasmids were designed to contain approximately 1000 bp up- and downstream regions around the deletion site, amplified from LAC* genomic DNA (Boles *et al*., 2010), and the first and last 30 bp of the open reading frame to be deleted.

For construction of plasmid pIMAY-Δ*957*, the up- and downstream regions of *SAUSA300_0957* were amplified using primers P2370/P2371 and P2372/P2373, spliced together in a second PCR using primers P2370/P2373 and cloned into pIMAY using XmaI/EcoRI. Plasmid pIMAY-Δ957 was recovered in *E. coli* XL1-Blue, creating strain ANG4147, shuttled through *E. coli* IM08B (ANG4159) and subsequently introduced into S. *aureus* LAC*Δ*957*. The allelic exchange to delete *SAUSA300_0957* and to create strain LAC*Δ*957* (ANG4290) was performed as previously described (Monk *et al*., 2012). For complementation analysis, plasmid piTET-*957* was integrated into the chromosome of strain LAC*Δ*957*, giving rise to the complementation strain LAC*Δ*957* piTET-*957* (ANG4341). As control, the empty plasmid piTET from ANG4163 was also integrated into the chromosome of LAC*Δ*957* yielding strain LAC*Δ*957* piTET (ANG4340). The *lcpB* (SAUSA300_0958) gene was deleted in a similar manner using primers P2844/P2845 and P2846/P2847 for the first and primers P2844/P2847 for the second PCR and cloning the fragment XmaI/EcoRI into pIMAY. Plasmid pIMAY-Δ*lcpB* was recovered in *E. coli* XL1blue (ANG4740), shuttled through *E. coli* IM08B (ANG4742) and introduced into the *S. aureus* LAC* (ANG4744) and finally yielding the *lcpB* deletion strain LAC*Δ*lcpB* (ANG4748).

Plasmid pIMAY*-Δ*957-958* was constructed for the production of an *SAUSA300_0957-0958* (*lcpB*) double mutant strain in which the first 30 bases of *SAUSA300_0957* were fused to the last 30 bp of *SAUSA300_0958* (*lcpB*). This was done using primers P2844/P2848 and P2849/P2373 in the first and primers P2844/P2373 in a second PCR. The fragment was cloned using XmaI/EcoRI into pIMAY* (Schuster *et al*., 2019) and the resulting plasmid pIMAY*-Δ*957-958* recovered in *E. coli* XL1-Blue (ANG4741). The plasmid was shuttled through *E. coli* IM08B (ANG4743), introduced into *S. aureus* LAC* (ANG4745) and following the allelic exchange procedure yielding the *S. aureus* strain LAC*Δ*957-958* (*lcpB*) (ANG4749).

The plasmid pIMAY-Δ*mgtE* was designed to contain up- and downstream regions of *SAUSA300_0910* (*mgtE*) fused together by the first and last 30 bp of the gene in order to create a LAC*Δ*mgtE* mutant. For this, primer pairs P2374/P2375 and P2376/P2377 were used to amplify up- and downstream regions and subsequently spliced together by PCR using primers P2374/P2377. The PCR fragment was digested with XmaI/EcoRI and ligated into pIMAY cut with the same enzymes. The plasmid pIMAY-Δ*mgtE* was recovered in *E. coli* XL1-Blue yielding strain ANG4146 and shuttled through *E. coli* IM08B yielding strain ANG4158. The plasmid pIMAY-Δ*mgtE* was then introduced into LAC* yielding strain LAC*Δ*SAUSA300_0910* (*mgtE*) (ANG4422) after the knockout procedure. To create an empty vector-containing control strain and a plasmid-based complementation strain, the plasmid piTET and piTET-*910* from strains ANG4163 and ANG4155 were isolated and used to transform strain LAC*Δ*SAUSA300_0910* (*mgtE*) (ANG4422), yielding strains ANG4445 (LAC*Δ*mgtE* piTET) and ANG4446 (LAC*Δ*mgtE* piTET-*mgtE*).

The *tagO* gene (*SAUSA300_0731*) was inactivated using the targetron intron homing system (Yao *et al*., 2006) or by allelic exchange using pIMAY* (Schuster *et al*., 2019). For the targetron mutagenesis, plasmid pNL9164-*tagO* was produced by amplifying the upstream targetron fragment with primers P2887 and the EBS universal primer (P2890) and the downstream fragment using primers P2888/P2889, the fragments were fused using the IBS primer (P2887) and the EBS1d primer (P2888). The final fragment was cloned using HindIII/BsrGI into pNL9164 (Yao *et al*., 2006). Plasmid pNL9164-*tagO* was recovered in *E. coli* XL1-Blue, yielding strain ANG4703, introduced into *E. coli* IM08B to give rise to strain ANG4704 and finally introduced into *S. aureus* LAC* (Boles *et al*., 2010) to create strain LAC* pNL9164-*tagO* (ANG4751). The final *tagO* targetron mutant strain LAC\**tagO*::targetron (ANG4753) was generated using the method described by Yao *et al*. (Yao *et al*., 2006). For the construction of *S. aureus* strain LAC*Δ*tagO* with an in-frame deletion in *tagO*, the allelic exchange plasmid pIMAY*-Δ*tagO* was used. To this end, up- and downstream *tagO* fragments were produced using primer pairs P2883/P2884 and P2885/P2886 and introduced by Gibson cloning into XmaI/EcoRI pre-cut pIMAY*. Plasmid pIMAY*-Δ*tagO* was recovered in *E. coli* XL1-Blue, generating strain ANG4755. After shuttling through *E. coli* strain IM08B (ANG4756), the plasmid was introduced into *S. aureus* LAC* to create strain ANG4757. Finally, the *tagO* locus was deleted by allelic exchange, generating strain LAC*Δ*tagO* (ANG4759). Because initial generation of the mutant failed, cells were plated unselectively onto TSA plates and incubated at room temperature (18-25°C) for about two weeks to enable colony differentiation between WT and Δ*tagO* mutant strains. Upon prolonged incubation, *tagO* mutant strain colonies have an opaque appearance.

## Data access

The whole genome sequencing data were deposited at the European Nucleotide Archive (ENA) under accession id ERP115099 and TN-seq data in the short read archive (SRA) at the National Center for Biotechnology Information (NCBI) under BioProject ID PRJNA544248. Analyzed tables can be found in the supplementary files. For other data, please contact the corresponding author.

## Acknowledgements

This research was funded by the Wellcome Trust through grants 100289 and 210671/Z/18/Z to A.G. and the NIH grant P01AI083214 to S.W.. C.F.S. was supported by the German Research Foundation [Deutsche Forschungsgemeinschaft (DFG)] through grant SCHU 3159/1–1 and F.C.M.K. by a Wellcome trust Ph.D. studentship. The funders had no role in study design, data collection and analysis, decision to publish, or preparation of the manuscript. We thank Microbes NG and the London Institute of Medical Sciences for providing whole genome sequencing services and the Tufts sequencing facility for the sequencing services for the TN-seq experiments. We also would like to thank Ute Bertsche for kind advice on HPLC analyses.

## Author contributions

C.F.S., D.M.W., F.C.K. and A.G. performed the research; C.F.S., D.M.W., F.C.M.K., M.S., S.W. and A.G. analyzed the data; C.F.S. and A.G. designed the research; C.F.S. and A.G. wrote the paper. All authors approved the final version of the manuscript.

## Disclosure declaration

The authors declare not conflict of interest.

## Graphical abstract

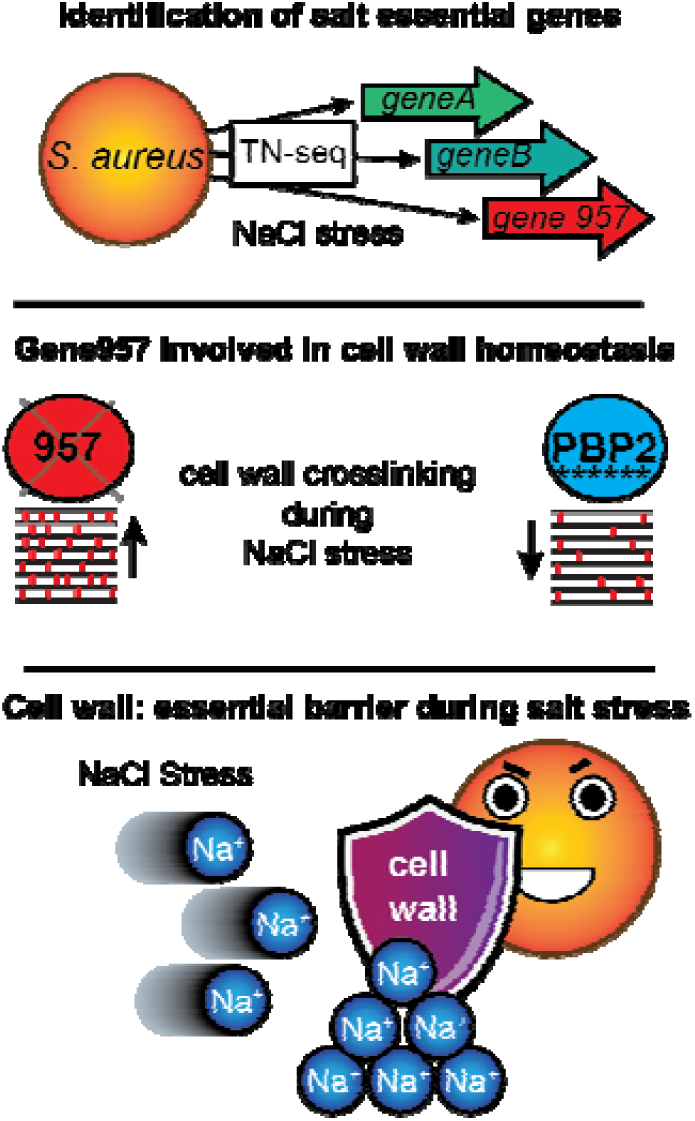

## Abbreviated summary

*Staphylococcus aureus* is able to grow in the presence of high concentrations of NaCl but the exact genetic factors contributing to this are unknown. Using a high-throughput TN-seq approach, we identified gene *957* as an important factor for the salt stress resistance in *S. aureus*. A *957*-mutant was not only salt sensitive but also showed increased peptidoglycan crosslinking, altogether highlighting the cell wall as an important barrier against osmotic stress.

## Notes

#### Summary of Updates

Data from sucrose and KCL Tn-seq experiments were again included from an earlier version

