## Supplementary material for "High-throughput transposon sequencing highlights the cell wall as an important barrier for osmotic stress in methicillin resistant *Staphylococcus aureus* and underlines a tailored response to different osmotic stressors": Table S15

**Schuster *et al.*, 2019**

**Table 15: Strains used**

| **Phages** | **Phage strain** |  | **Origin** |
| --- | --- | --- | --- |
|  | Phage Φ85 |  | Olaf Schneewind lab (Chicago) |
|  | Phage Φ11-FRT |  | (Santiago *et al.* 2015) |
| ***E. coli*** | **Strain** | **Resistance** | **Reference** |
|  | XL1-Blue cloning strain (ANG127) | Tet^R^ | Stratagene |
|  | XL1-Blue pCL55 (ANG243) | Amp^R^ | (Lee *et al.* 1991) |
|  | XL1-Blue piTET (ANG284) | Amp^R^ | (Gründling and Schneewind 2007) |
|  | XL1-Blue pNL9164 (ANG1432) | Amp^R^ | Sigma Aldrich |
|  | DH10B pIMAY (ANG2154) | Cm^R^ | (Monk *et al.* 2012) |
|  | IM08B WT (ANG3724) |  | (Monk *et al.* 2015) |
| ANG3732 | IM08B pCL55 | Amp^R^ | (Schuster *et al.* 2016) |
| ANG3928/4163 | IM08B piTET | Amp^R^ | (Karinou *et al.* 2019) |
| ANG4005 | XL1-Blue pIMAY* | Cm^R^ | (Schuster *et al.* 2019) |
| ANG4139 | XL1-Blue piTET-*SAUSA300_0481* | Amp^R^ | This study |
| ANG4140 | XL1-Blue piTET-*SAUSA300_0482* | Amp^R^ | This study |
| ANG4142 | XL1-Blue piTET-*SAUSA300_0694* | Amp^R^ | This study |
| ANG4144 | XL1-Blue piTET-*SAUSA300_0910* | Amp^R^ | This study |
| ANG4145 | XL1-Blue piTET-*SAUSA300_0957* | Amp^R^ | This study |
| ANG4146 | XL1-Blue pIMAY-∆*SAUSA300_0910 (mgtE)* | Cm^R^ | This study |
| ANG4147 | XL1-Blue pIMAY-∆*SAUSA300_0957* | Cm^R^ | This study |
| ANG4150 | IM08B piTET-*SAUSA300_0481* | Amp^R^ | This study |
| ANG4151 | IM08B piTET-*SAUSA300_0482* | Amp^R^ | This study |
| ANG4153 | IM08B piTET-*SAUSA300_0694* | Amp^R^ | This study |
| ANG4156 | IM08B piTET-*SAUSA300_0957* | Amp^R^ | This study |
| ANG4158 | IM08B piTET-*SAUSA300_0910* | Amp^R^ | This study |
| ANG4158 | IM08B pIMAY-∆*SAUSA300_0910 (mgtE)* | Cm^R^ | This study |
| ANG4159 | IM08B pIMAY-∆*SAUSA300_0957* | Cm^R^ | This study |
| ANG4291 | XL1-Blue pCL55-*SAUSA300_0483* | Amp^R^ | This study |
| ANG4292 | IM08B pCL55-*SAUSA300_0483* | Amp^R^ | This study |
| ANG4293 | XL1-Blue pCL55-*SAUSA300_0867* | Amp^R^ | This study |
| ANG4294 | IM08B pCL55-*SAUSA300_0867* | Amp^R^ | This study |
| ANG4703 | XL1-Blue pNL9164-*tagO* | Amp^R^ | This study |
| ANG4704 | IM08B pNL9164-*tagO* | Amp^R^ | This study |
| ANG4740 | XL1-Blue pIMAY*-∆*SAUSA300_0958 (lcpB)* | Cm^R^ | This study |
| ANG4741 | XL1-Blue pIMAY*-∆*SAUSA300_0958-(lcpB)*∆*SAUSA300_0957* | Cm^R^ | This study |
| ANG4742 | IM08B pIMAY*-∆*SAUSA300_0958 (lcpB)* | Cm^R^ | This study |
| ANG4743 | IM08B pIMAY*-∆*SAUSA300_0958-(lcpB)*∆*SAUSA300_0957* | Cm^R^ | This study |
| ANG4755 | XL1-Blue pIMAY*-∆*tagO* | Cm^R^ | This study |
| ANG4756 | IM08B pIMAY*-∆*tagO* | Cm^R^ | This study |
| **Strain number** | **Strain name** | **Resistance** | **Reference** |
| AH1263 | LAC* (ANG1575) |  | (Boles *et al.* 2010) |
|  | JE2 (ANG2624) |  | (Fey *et al.* 2013) |
|  | TM283 (USA300 TCH1516 without pUSA300HUMR; ANG3729) |  | (Coe *et al.* 2019) |
|  | TM283 pORF5-Tn(+) (ANG3801) | Kan^R^, 30°C | (Coe *et al.* 2019) |
|  | TM283 pORF5-Tn(-) (ANG3803) | Kan^R^, 30°C | (Coe *et al.* 2019) |
| NE157 | JE2 *SAUSA300_0678::Tn* (ANG3901) | Erm^R^ | (Fey *et al.* 2013) |
| NE526 | JE2 *SAUSA300_0694::Tn* (ANG3902) | Erm^R^ | (Fey *et al.* 2013) |
| NE736 | JE2 *SAUSA300_0910::Tn* (ANG3904) | Erm^R^ | (Fey *et al.* 2013) |
| NE810 | JE2 *SAUSA300_1642::Tn* (ANG3906) | Erm^R^ | (Fey *et al.* 2013) |
| NE997 | JE2 *SAUSA300_2071::Tn* (ANG3908) | Erm^R^ | (Fey *et al.* 2013) |
| NE1384 | JE2 SAUSA300*_0957::Tn* (ANG3911) | Erm^R^ | (Fey *et al.* 2013) |
| NE1458 | JE2 *SAUSA300_2094::Tn* (ANG3912) | Erm^R^ | (Fey *et al.* 2013) |
| NE779 | JE2 *SAUSA300_1254::Tn* (ANG3929) | Erm^R^ | (Fey *et al.* 2013) |
| NE188 | JE2 *SAUSA300_0481::Tn* (ANG3968) | Erm^R^ | (Fey *et al.* 2013) |
| NE251 | JE2 *SAUSA300_0482::Tn* (ANG3969) | Erm^R^ | (Fey *et al.* 2013) |
| NE431 | JE2 *SAUSA300_0621::Tn* (ANG3970) | Erm^R^ | (Fey *et al.* 2013) |
| NE629 | JE2 *SAUSA300_0726::Tn* (ANG3971) | Erm^R^ | (Fey *et al.* 2013) |
| NE723 | JE2 *SAUSA300_1514::Tn* (ANG3972) | Erm^R^ | (Fey *et al.* 2013) |
| NE867 | JE2 *SAUSA300_0483::Tn* (ANG3974) | Erm^R^ | (Fey *et al.* 2013) |
| NE987 | JE2 *SAUSA300_2249::Tn* (ANG3975) | Erm^R^ | (Fey *et al.* 2013) |
| NE1109 | JE2 *SAUSA300_2022::Tn* (ANG3976) | Erm^R^ | (Fey *et al.* 2013) |
| NE1340 | JE2 *SAUSA300_1692::Tn* (ANG3978) | Erm^R^ | (Fey *et al.* 2013) |
| NE1472 | JE2 *SAUSA300_2023::Tn* (ANG3979) | Erm^R^ | (Fey *et al.* 2013) |
| NE1494 | JE2 *SAUSA300_1126::Tn* (ANG3980) | Erm^R^ | (Fey *et al.* 2013) |
| NE1778 | JE2 *SAUSA300_0958::Tn* (ANG3981) | Erm^R^ | (Fey *et al.* 2013) |
| NE1833 | JE2 *SAUSA300_2026::Tn* (ANG3982) | Erm^R^ | (Fey *et al.* 2013) |
| NE1841 | JE2 *SAUSA300_1474::Tn* (ANG3983) | Erm^R^ | (Fey *et al.* 2013) |
| NE1872 | JE2 *SAUSA300_2024::Tn* (ANG3984) | Erm^R^ | (Fey *et al.* 2013) |
| NE535 | JE2 *SAUSA300_0867::Tn* (ANG3986) | Erm^R^ | (Fey *et al.* 2013) |
| ANG4054 | LAC* piTET | Cm^R^ | (Zeden *et al.* 2018) |
| ANG4290 | LAC* ∆*SAUSA300_0957* |  | This study |
| ANG4307 | JE2 piTET | Cm^R^ | This study |
| ANG4308 | NE188 *SAUSA300_0481::Tn* piTET | Erm^R^/Cm^R^ | This study |
| ANG4309 | NE188 *SAUSA300_0481::Tn* piTET-*SAUSA300_0481* | Erm^R^/Cm^R^ | This study |
| ANG4310 | NE251 *SAUSA300_0482::Tn* piTET | Erm^R^/Cm^R^ | This study |
| ANG4311 | NE251 *SAUSA300_0482::Tn* piTET-*SAUSA300_0482* | Erm^R^/Cm^R^ | This study |
| ANG4325 | JE2 pCL55 | Cm^R^ | This study |
| ANG4326 | NE867 *SAUSA300_0483::Tn* pCL55 | Erm^R^/Cm^R^ | This study |
| ANG4327 | NE867 *SAUSA300_0483::Tn* pCL55-*SAUSA300_0483* | Erm^R^/Cm^R^ | This study |
| ANG4328 | NE535 *SAUSA300_0867::Tn* pCL55 | Erm^R^/Cm^R^ | This study |
| ANG4329 | NE535 *SAUSA300_0867::Tn* pCL55-*SAUSA300_0867* | Erm^R^/Cm^R^ | This study |
| ANG4336 | NE736 *SAUSA300_0910::Tn* piTET | Erm^R^/Cm^R^ | This study |
| ANG4337 | NE736 *SAUSA300_0910::Tn* piTET-*SAUSA300_0910* | Erm^R^/Cm^R^ | This study |
| ANG4338 | NE1384 *SAUSA300_0957::Tn* piTET | Erm^R^/Cm^R^ | This study |
| ANG4339 | NE1384 *SAUSA300_0957::Tn* piTET-*SAUSA300_0957* | Erm^R^/Cm^R^ | This study |
| ANG4340 | LAC* ∆*SAUSA300_0957* piTET | Cm^R^ | This study |
| ANG4341 | LAC* ∆*SAUSA300_0957* piTET-*SAUSA300_0957* | Cm^R^ | This study |
| ANG4377 | NE526 *SAUSA300_0694::Tn* piTET | Erm^R^/Cm^R^ | This study |
| ANG4378 | NE526 *SAUSA300_0694::Tn* piTET-*SAUSA300_0694* | Erm^R^/Cm^R^ | This study |
| ANG4381 | LAC* ∆*SAUSA300_0957* Suppressor S1 |  | This study |
| ANG4382 | LAC* ∆*SAUSA300_0957* Suppressor S2 |  | This study |
| ANG4383 | LAC* ∆*SAUSA300_0957* Suppressor S3 |  | This study |
| ANG4384 | LAC* ∆*SAUSA300_0957* Suppressor S4 |  | This study |
| ANG4386 | LAC* ∆*SAUSA300_0957* Suppressor S5 |  | This study |
| ANG4389 | LAC* ∆*SAUSA300_0957* Suppressor S6 |  | This study |
| ANG4390 | LAC* ∆*SAUSA300_0957* Suppressor S7 |  | This study |
| ANG4391 | LAC* ∆*SAUSA300_0957* Suppressor S8 |  | This study |
| ANG4393 | LAC* ∆*SAUSA300_0957* Suppressor S9 |  | This study |
| ANG4394 | LAC* ∆*SAUSA300_0957* Suppressor S10 |  | This study |
| ANG4422 | LAC* ∆*SAUSA300_0910* (*mgtE*) |  | This study |
| ANG4445 | LAC* ∆*SAUSA300_0910* (*mgtE*) piTET | Cm^R^ | This study |
| ANG4446 | LAC* ∆*SAUSA300_0910* (*mgtE*) piTET-*SAUSA300_0957* | Cm^R^ | This study |
| ANG4485 | NE789 SAUSA300_1332::Tn | Erm^R^ | This study |
| ANG4527 | LAC* ∆*SAUSA300_0957* Suppressor S2 *SAUSA300_1332::Tn* | Erm^R^ | This study |
| ANG4528 | LAC* ∆*SAUSA300_0957* Suppressor S4 *SAUSA300_1332::Tn* | Erm^R^ | This study |
| ANG4557 | LAC* ∆*SAUSA300_0957* Suppressor S2 *SAUSA300_1332::Tn* repaired (WT) *pbp2* | Erm^R^ | This study |
| ANG4558 | LAC* ∆*SAUSA300_0957* Suppressor S4 *SAUSA300_1332::Tn* repaired (WT) *pbp2* | Erm^R^ | This study |
| ANG4561 | LAC* *SAUSA300_1332::Tn* | Erm^R^ | This study |
| ANG4562 | LAC* ∆*SAUSA300_0957* *SAUSA300_1332::Tn* | Erm^R^ | This study |
| ANG4563 | LAC* *SAUSA300_1332::Tn* *pbp2* SNP S2 | Erm^R^ | This study |
| ANG4564 | LAC* *SAUSA300_1332::Tn* *pbp2* SNP S4 | Erm^R^ | This study |
| ANG4624 | LAC* ∆*SAUSA300_0957 SAUSA300_1332::Tn pbp2* SNP S2 | Erm^R^ | This study |
| ANG4625 | LAC* ∆*SAUSA300_0957* *SAUSA300_1332::Tn* *pbp2* SNP S4 | Erm^R^ | This study |
| ANG4744 | LAC* pIMAY*-∆*SAUSA300_0958* (*lcpB*) | Cm^R^, 28°C | This study |
| ANG4745 | LAC* pIMAY-∆*SAUSA300_0958*-(*lcpB*)∆*SAUSA300_0957* | Cm^R^, 28°C | This study |
| ANG4748 | LAC* ∆*SAUSA300_0958* (*lcpB*) |  | This study |
| ANG4749 | LAC* ∆*SAUSA300_0958* (*lcpB*)∆*SAUSA300_0957* |  | This study |
| ANG4751 | LAC* pNL9164-*tagO* | Erm^R^ | This study |
| ANG4753 | LAC* LAC* *tagO::targetron* | Erm^S^ | This study |
| ANG4757 | LAC* pIMAY*-∆*tagO* | Cm^R^, 28°C | This study |
| ANG4759 | LAC* ∆*tagO* |  | This study |

Boles BR, Thoendel M, Roth AJ, Horswill AR. 2010. Identification of genes involved in polysaccharide-independent *Staphylococcus aureus* biofilm formation. *PLoS One* **5**: e10146.

Coe KA, Lee W, Komazin-Meredith G, Meredith TC, Grad YH, Walker S. 2019. Comparative Tn-Seq reveals common daptomycin resistance determinants in *Staphylococcus aureus* despite strain-dependent differences in essentiality of shared cell envelope genes. *bioRxiv* doi:10.1101/648246: 648246.

Fey PD, Endres JL, Yajjala VK, Widhelm TJ, Boissy RJ, Bose JL, Bayles KW. 2013. A genetic resource for rapid and comprehensive phenotype screening of nonessential *Staphylococcus aureus* genes. *MBio* **4**: e00537-00512.

Gründling A, Schneewind O. 2007. Genes required for glycolipid synthesis and lipoteichoic acid anchoring in *Staphylococcus aureus*. *J Bacteriol* **189**: 2521-2530.

Karinou E, Schuster CF, Pazos M, Vollmer W, Gründling A. 2019. Inactivation of the Monofunctional Peptidoglycan Glycosyltransferase SgtB Allows *Staphylococcus aureus* To Survive in the Absence of Lipoteichoic Acid. *J Bacteriol* **201**.

Lee CY, Buranen SL, Ye ZH. 1991. Construction of single-copy integration vectors for *Staphylococcus aureus*. *Gene* **103**: 101-105.

Monk IR, Shah IM, Xu M, Tan MW, Foster TJ. 2012. Transforming the untransformable: application of direct transformation to manipulate genetically *Staphylococcus* *aureus* and *Staphylococcus epidermidis*. *MBio* **3**.

Monk IR, Tree JJ, Howden BP, Stinear TP, Foster TJ. 2015. Complete Bypass of Restriction Systems for Major *Staphylococcus aureus* Lineages. *MBio* **6**: e00308-00315.

Santiago M, Matano LM, Moussa SH, Gilmore MS, Walker S, Meredith TC. 2015. A new platform for ultra-high density *Staphylococcus aureus* transposon libraries. *BMC Genomics* **16**: 252.

Schuster CF, Bellows LE, Tosi T, Campeotto I, Corrigan RM, Freemont P, Gründling A. 2016. The second messenger c-di-AMP inhibits the osmolyte uptake system OpuC in *Staphylococcus aureus*. *Sci Signal* **9**: ra81.

Schuster CF, Howard SA, Gründling A. 2019. Use of the counter selectable marker PheS* for genome engineering in *Staphylococcus aureus*. *Microbiology* doi:10.1099/mic.0.000791.

Zeden MS, Schuster CF, Bowman L, Zhong Q, Williams HD, Gründling A. 2018. Cyclic di-adenosine monophosphate (c-di-AMP) is required for osmotic regulation in *Staphylococcus aureus* but dispensable for viability in anaerobic conditions. *J Biol Chem* **293**: 3180-3200.
