## Supplemental Data 1 for "High-throughput transposon sequencing highlights the cell wall as an important barrier for osmotic stress in methicillin resistant *Staphylococcus aureus* and underlines a tailored response to different osmotic stressors"

**Running title**: *Global analysis of osmotic stress responses in Staphylococcus aureus*

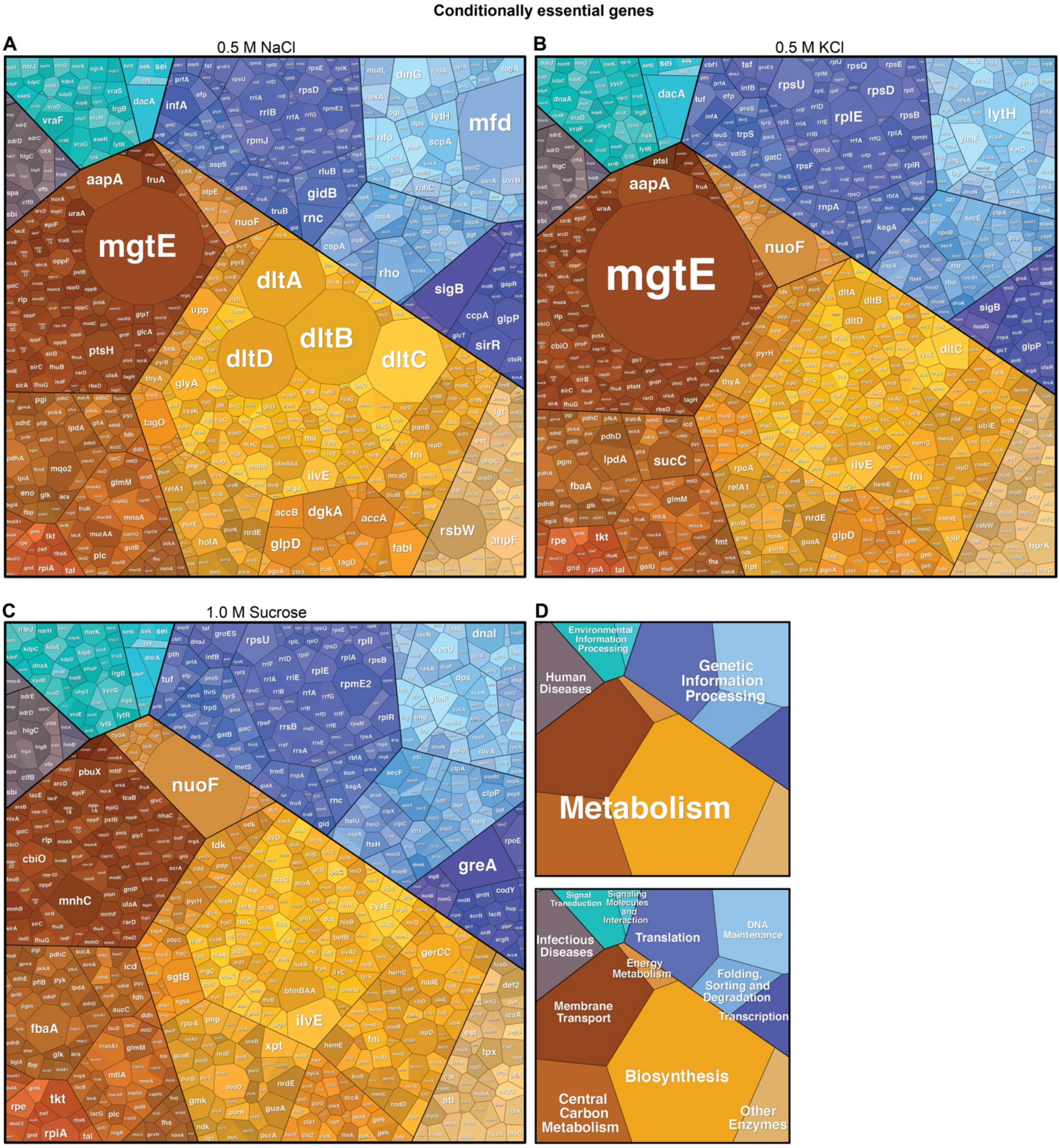


**Supplementary Fig. S1. Voronoi diagram-based visualization of conditionally essential *S. aureus* genes when exposed to different osmotic stressors underlines differences.**

A.-C. Conditionally essential genes of *S. aureus* strains grown either in 0.5 M NaCl (A), 0.5 M KCl (B) or 1.0 M sucrose (C). Area sizes were adjusted to their essentiality regardless of p- or q-value to visualize differences on a genome wide level. The larger the area, the higher the number of transposon insertions in the LB control condition as compared to the stress conditions. D. Overview of polygon assignment: colors of polygons indicate cellular functions.

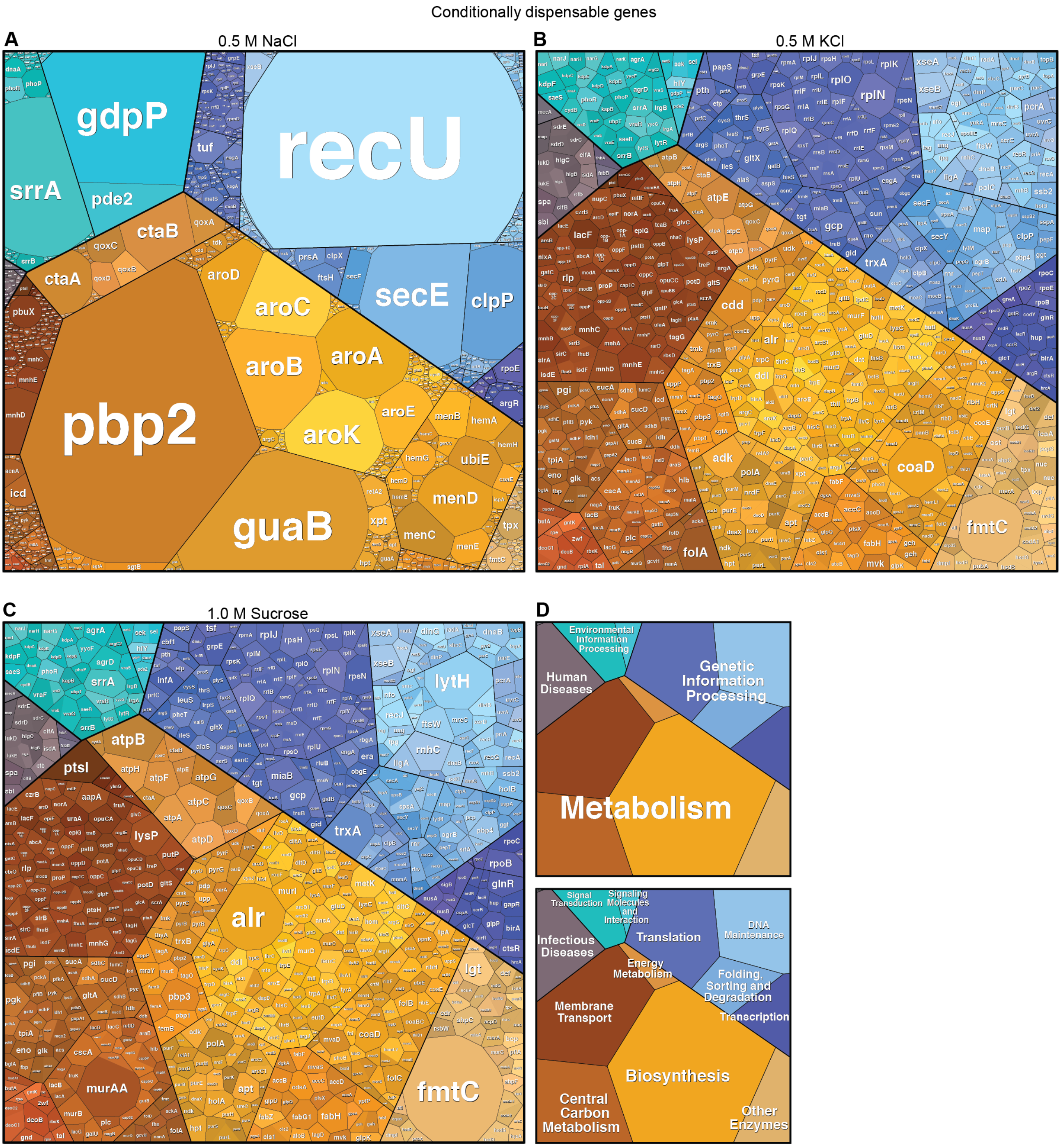


**Supplementary Fig. S2. Voronoi diagram-based visualization of conditionally dispensable *S. aureus* genes when exposed to different osmotic stressors.**

A.-C. Conditionally dispensable genes of *S. aureus* strains grown either in 0.5 M NaCl (A), 0.5 M KCl (B) or 1.0 M sucrose (C) were mapped to cellular functions with area sizes adjusted to their essentiality regardless of p- or q-value. The larger the area, the higher the number of transposon insertions in relation to the LB control. *recU* was scaled to 1/10 of the original ratio to preserve space. D. Overview of polygon assignment: colors of polygons indicate cellular functions.

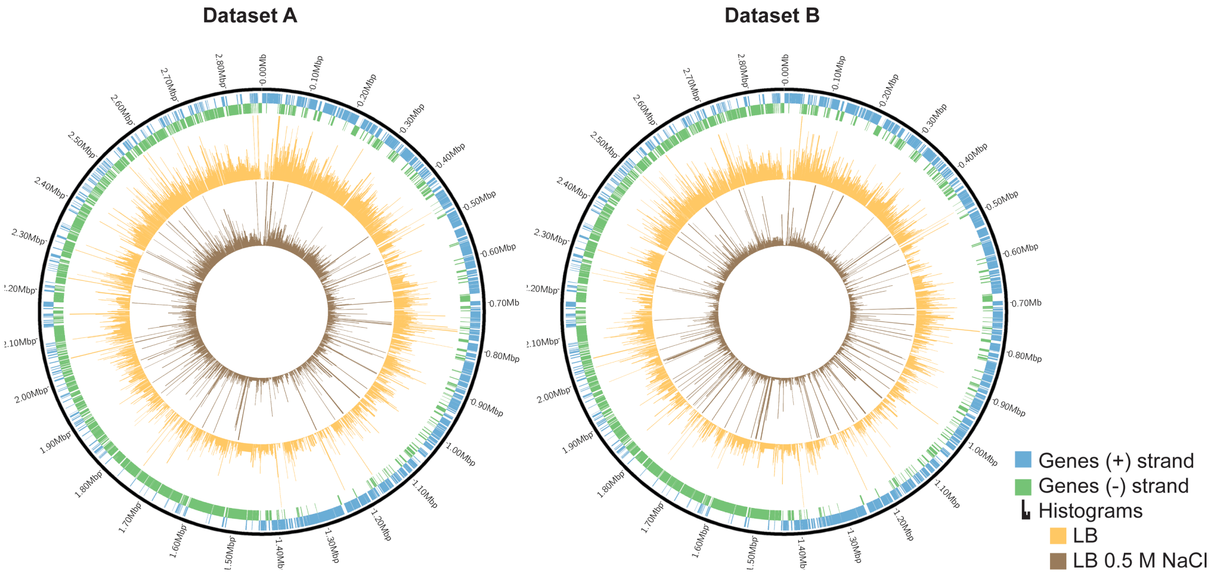


**Supplementary Fig. S3. Circos plots showing transposon insertions in the *S. aureus* genome following growth under different conditions.**

The outer two bands represent genes on the forward (blue) and reverse (green) strand, respectively. The inner two rings represent transposon insertions per gene following the growth of *S. aureus* in in either LB (orange) or LB 0.5 M NaCl (brown) for 17 generations. Each circular plot represents one individual experiment (n=1).

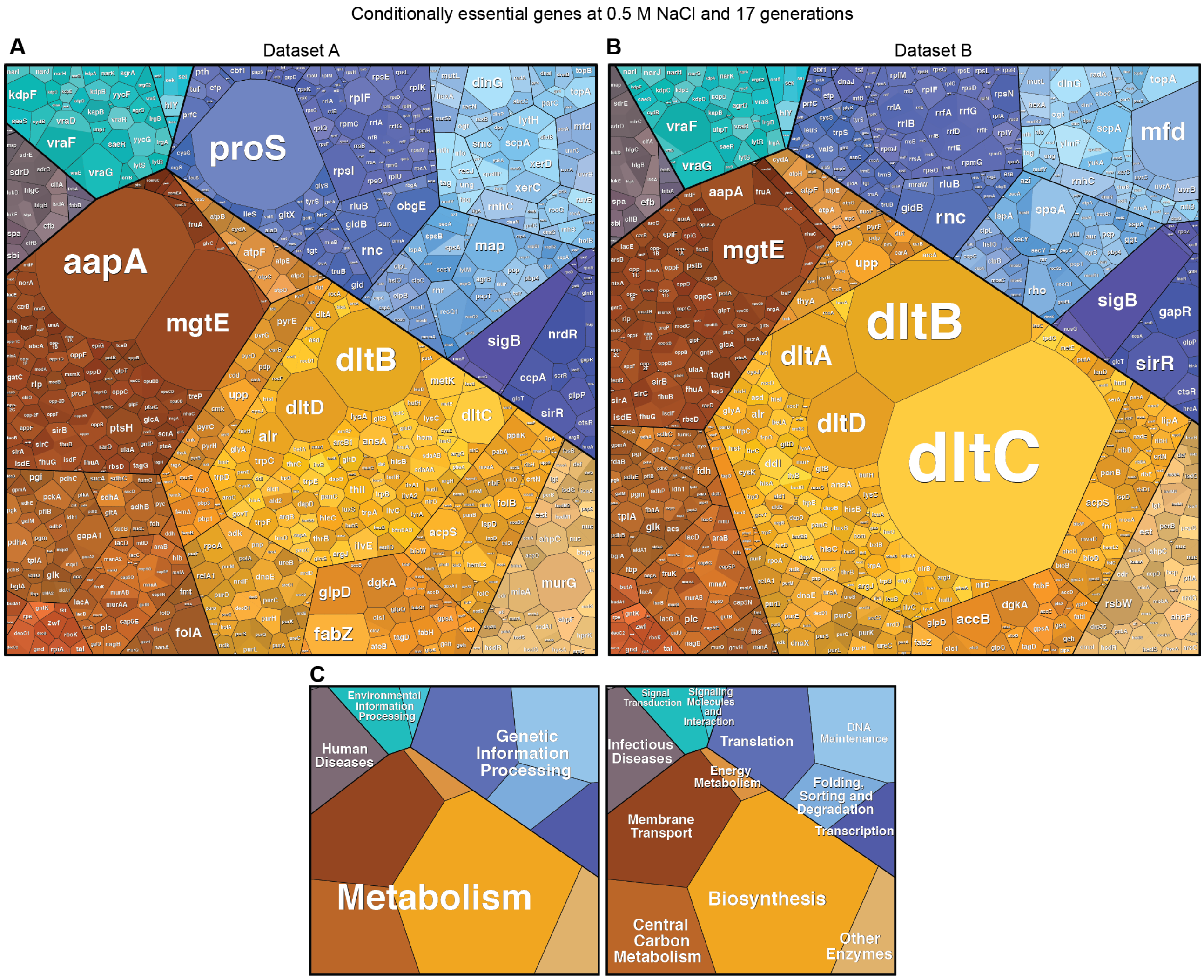


**Supplementary Fig. S4. Voronoi diagram-based visualization of *S. aureus* conditionally essential genes when exposed to 0.5 M NaCl.**

A.-B. Conditionally essential *S. aureus* genes from replicate A (A) and replicate B (B) grown in 0.5 M NaCl for 17 generations were mapped to cellular functions with area sizes adjusted to their essentiality regardless of p- or q-value. The larger the area, the lower the number of transposon insertions in stress condition as compared to the LB control condition. C. Overview of polygon assignment: colors of polygons indicate cellular functions.

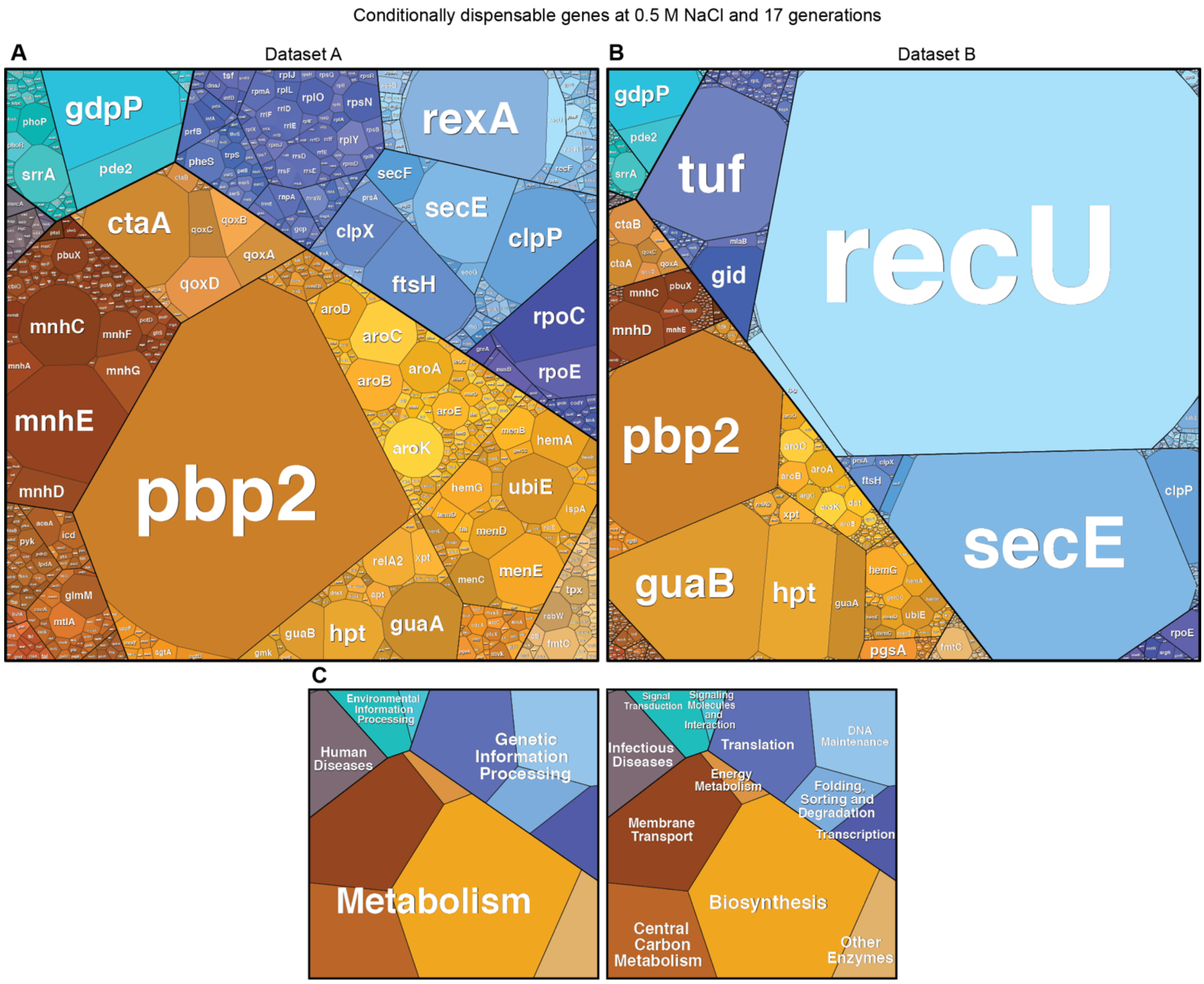


**Supplementary Fig. S5. Voronoi diagram-based visualization of *S. aureus* conditionally dispensable genes when exposed to 0.5 M NaCl.**

A.-B. Conditionally dispensable *S. aureus* genes from replicate A (A) and replicate B (B) grown in 0.5 M NaCl for 17 generations were mapped to cellular functions with area sizes adjusted to their dispensability regardless of p- or q-value. The larger the area, the higher the number of transposon insertions in the stress condition as compared to the LB control. *recU* was scaled to 1/5 of the original ratio to preserve space. C. Overview of polygon assignment: colors of polygons indicate cellular functions.

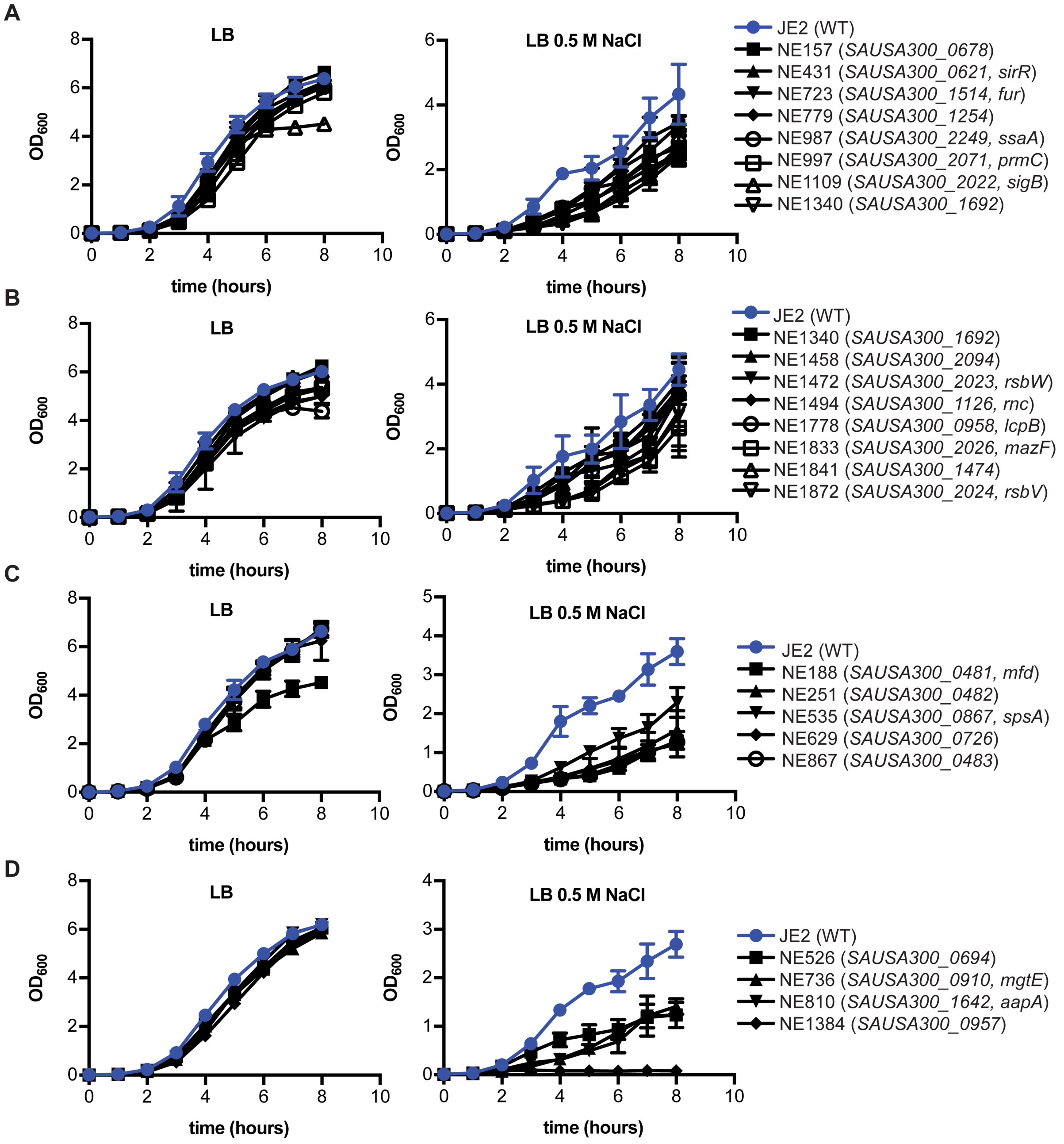


**Supplementary Fig. S6. Growth curves of *S. aureus* strains with transposon insertions in potential salt essential genes.**

A-D. *S. aureus* strain JE2 (WT) and strains from the Nebraska Transposon Mutant Library (NTML) containing transposon insertions in potential salt essential genes were grown in LB (left column) or 0.5 M NaCl LB medium (right column) and their growth monitored over 8 hours. Transposon mutant strains shown in panels A and B had a mild growth defect while strains shown in panels C and D had a stronger growth defect when grown in the high salt medium. NE numbers correspond to the NTML mutant number. Locus tag number and where available gene names with transposon insertion site are given in parenthesis. Growth curves were performed in triplicates (n=3) and means and SDs were plotted.

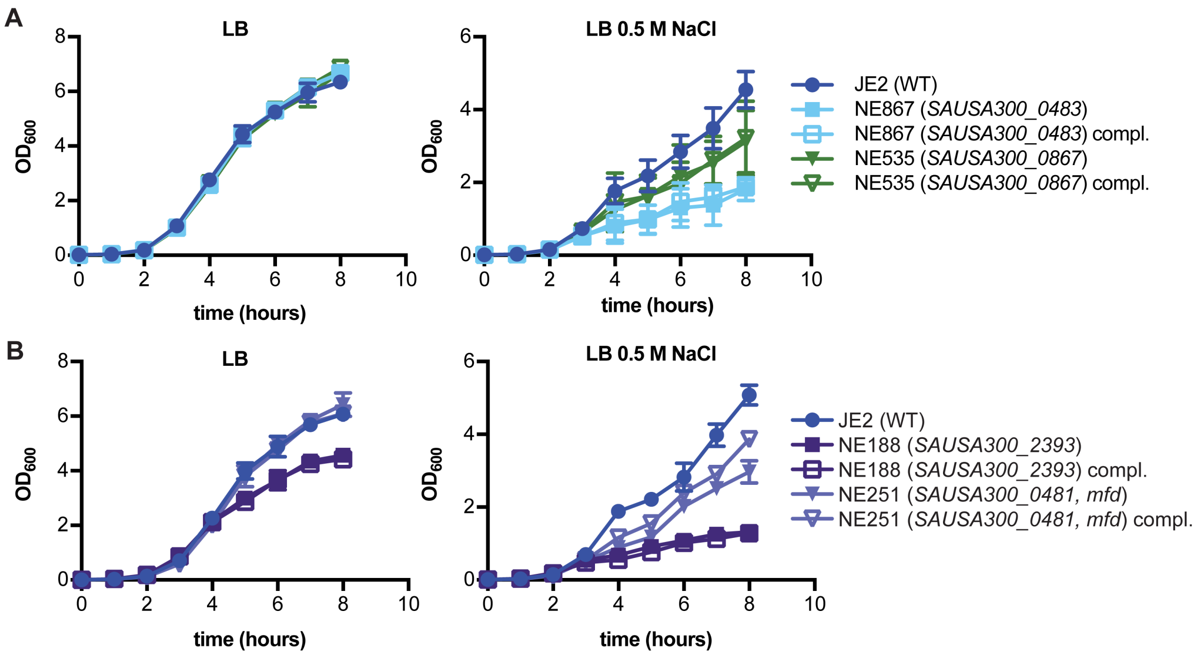


**Supplementary Fig. S7. Complementation analysis using *S. aureus* mutants with transposon insertions in putative salt resistance genes.**

A. Growth curves. *S. aureus* strains JE2 (WT), NE867 and NE535 with integrated pCL55 and *S. aureus* strains NE867 and NE535 with respective complementation plasmids were grown in either LB or LB 0.5 M NaCl and their growth monitored by determining $OD_{600}$ readings.
B. As in panel A but using strains JE2 piTET (WT) or NE188 and NE251 containing plasmid piTET or the respective complementation plasmid. Strains were grown in LB or 0.5 M NaCl LB medium supplemented with 100 ng/ml Atet. All experiments were conducted three times and the means and standard deviations were plotted.

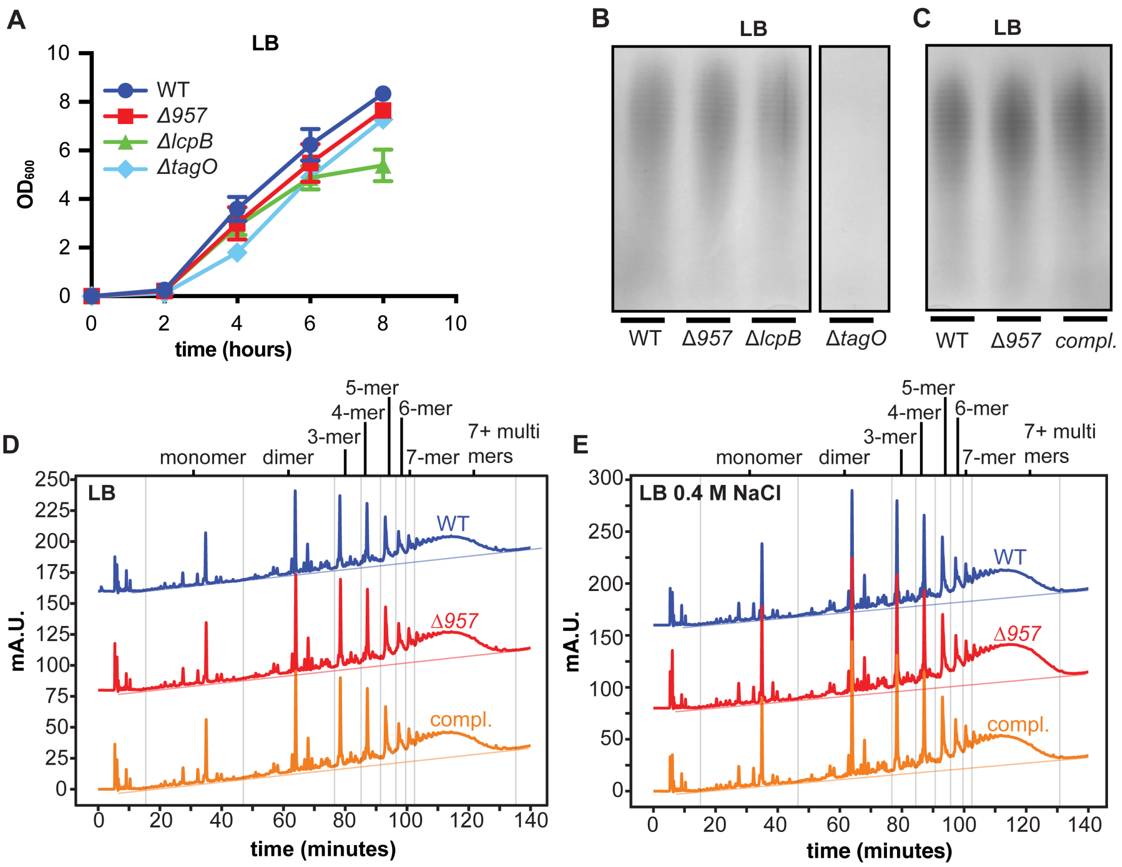


**Supplementary Fig. S8. Investigating the involvement of gene *957* in cell wall homeostasis.**

A. Growth curves using strains with mutations in WTA synthesis genes. *S. aureus* strains LAC* (WT), LAC*Δ*957* (Δ*957*), LAC*Δ*lcpB* (Δ*lcpB*) and LAC*Δ*tagO* (Δ*tagO*) were grown in LB and growth monitored by measuring $OD_{600}$ readings. The experiment was performed three times and means and standard deviations were plotted.

B. Detection of WTA. WTA was isolated from the strains described in panel A following growth in LB medium, then subjected to electrophoresis on polyacrylamide gels and visualized by silver staining. One representative result is shown out of three independent experiments.

C. Detection of WTA. Experimental setup is the same as in panel B but using the strains LAC* piTET (WT), LAC*Δ*957* piTET (Δ*957*) and LAC* Δ*957* piTET-*957* (compl.).

D. Muropeptide profile. The same strains as described in panel C were grown in LB medium and the peptidoglycan isolated, digestion with mutanolysin and the muropeptides subsequently separated by HPLC. Shown are representative chromatograms from three independent experiments. Retention time ranges of mono-, di- and higher multi-mers are indicated by vertical lines.

E. Muropeptide profiles. As in panel D but bacteria were grown in LB 0.4 M NaCl.

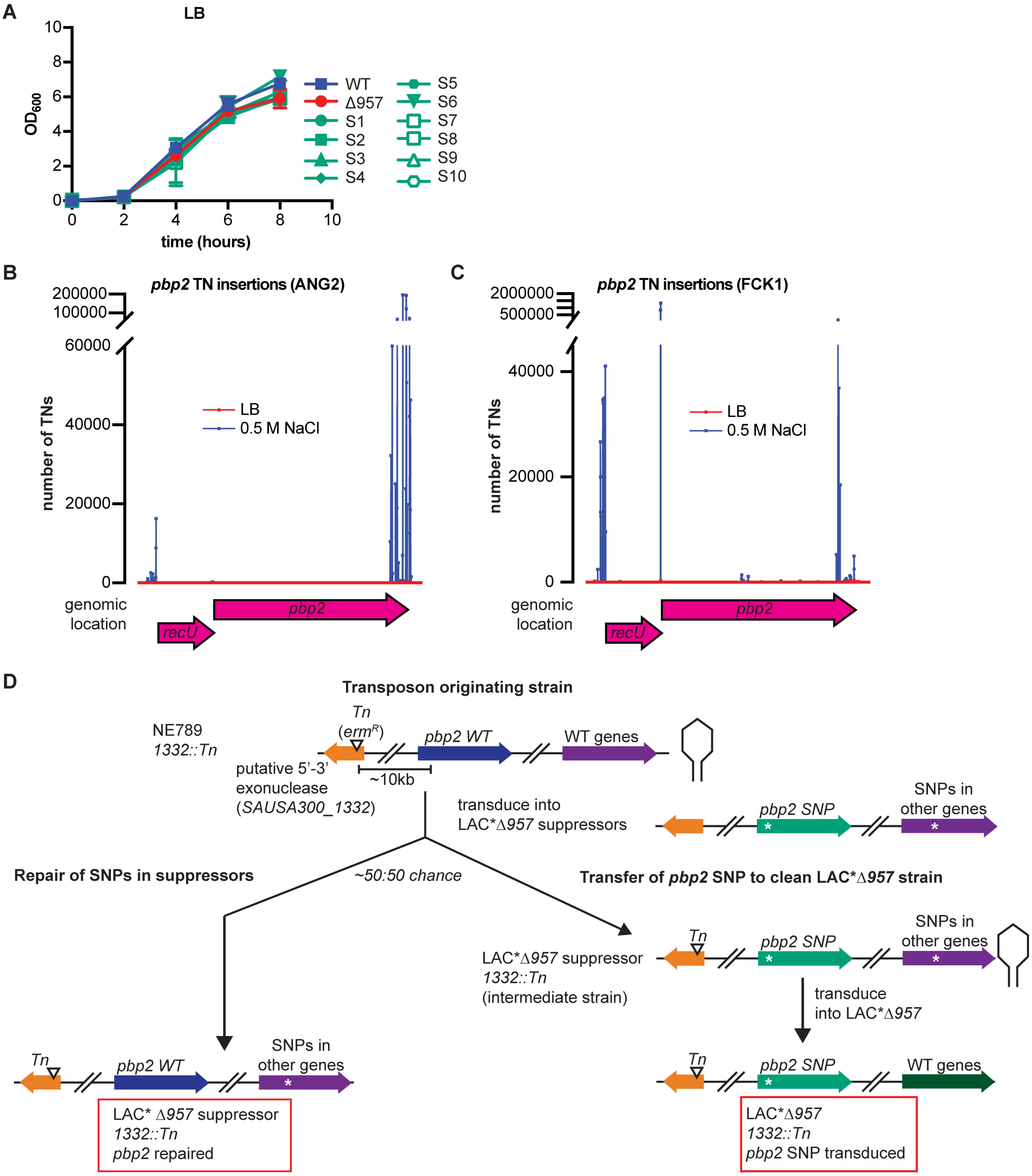


**Supplementary Fig. S9. Growth of *Δ957* suppressor strains and schematic of the co-transduction strategy for complementation analysis.**

A. Growth curves of *Δ957* suppressor strains. Strains LAC* (WT), LAC*Δ*957* (Δ*957*) and LAC*Δ*957* suppressors S1 trough S10 (S1-S10) were grown in LB medium and their growth monitored by taking $OD_{600}$ readings. The experiment was performed three times and means and standard deviations were plotted.

B. TN insertion map. Transposon insertions from replicate A (ANG dataset) were mapped to the *recU*/*pbp2* operon from a library grown in LB (red) or LB medium containing 0.5 M NaCl salt (blue).

C. TN insertion map. As in panel B but using replicate B (FCK dataset).

D. Schematic of phage co-transduction experiment to either repair the *pbp2* SNP in the suppressor strains (left branch) or to transduce the *pbp2* suppressor mutations into a clean *957* mutant (right branch). A phage lysate was prepared using an NTML library strain with a transposon insertion in an unrelated gene (*SAUSA300_1332*) located approximately 10 kb from the *pbp2* SNPs present in the suppressor strains. This lysate was used for transductions using different LAC*Δ*957* suppressor strains as recipient strains. With about 50% frequency, the *pbp2* SNPs were exchanged to the wild type allele by transducing *SAUSA300_1332*::*Tn* and the WT *pbp2* allele producing strain LAC*Δ*957* suppressor *1332*::*Tn pbp2* repaired (left branch). In the other cases, the *SAUSA300_1332*::*Tn* region was transferred to a suppressor strain without replacing the *pbp2* SNP (right branch). A phage lysate prepared on the LAC*Δ*957* suppressor *1332*::*Tn* intermediate strain was then used to transduce the *pbp2* suppressor SNP into a clean LAC*Δ*957* strain, resulting in the clean suppressor strain LAC*Δ*957 1332*::Tn *pbp2* SNP transduced.

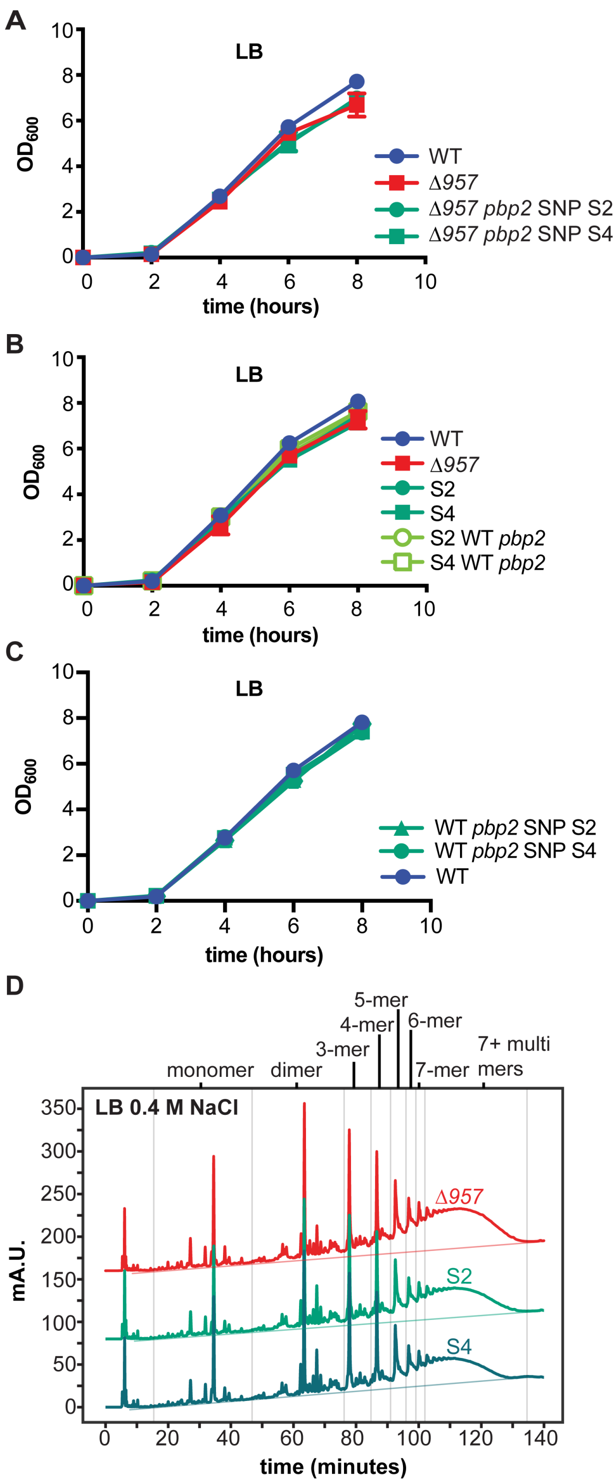


**Supplementary Fig. S10. Growth and peptidoglycan analysis of *957* mutant and suppressor strains.**

A. Growth curves using *957* mutants with transduced *pbp2* SNPs. Strains LAC* *1332*::Tn (WT), LAC*Δ*957 1332*::Tn (Δ*957*), LAC*Δ*957 1332*::Tn *pbp2* SNP S2 (Δ*957 pbp2* SNP S2), LAC*Δ*957* suppressor S4 *1332*::Tn (Δ*957 pbp2* SNP S4) were grown in LB medium and growth monitored by determining $OD_{600}$ readings. The means and standard deviations from three independent experiments were plotted.

B. Growth curves using *957* suppressor strains carrying a repaired WT *pbp2* gene. Same as in panel A but using strains LAC* *1332*::Tn (WT), LAC*Δ*957* *1332*::Tn (Δ*957*), LAC*Δ*957* suppressor S2 *1332*::Tn (S2), LAC*Δ*957* suppressor S4 *1332*::Tn (S4), LAC*Δ*957* suppressor S2 *1332*::Tn repaired WT *pbp2* (S2 WT *pbp2*), and LAC*Δ*957* suppressor S4 *1332*::Tn repaired WT *pbp2* (S4 WT *pbp2*).
C. Growth curves using WT strains carrying *pbp2* SNP mutations. Same as in panel A but using strains LAC* *1332*::Tn (WT), LAC* *1332*::Tn *pbp2* SNP S2 (WT *pbp2* SNP S2) and LAC* *1332*::Tn *pbp2* SNP S4 (WT *pbp2* SNP S4).
D. Muropeptide profiles. Strains LAC*Δ*957* (Δ*957*), LAC*Δ*957* Suppressor S2 (S2) and S4 LAC*Δ*957* Suppressor S4 (S4) were grown in LB medium with 0.4 M NaCl, the peptidoglycan was extracted, digested with mutanolysin, and the muropeptide separated by HPLC. Retention ranges for monomers, di- and higher oligomers are indicated by vertical lines. The experiment was conducted three times and one representative result is shown.
